# The *Drosophila* SPECC1L homolog, Split Discs, co-localizes with non-muscle myosin II and regulates focal adhesion dynamics

**DOI:** 10.1101/2025.04.06.647460

**Authors:** Aidan Teran, Jeehoon Jung, Amy Platenkamp, Alesandra Pardini, Andy Zhao, Sushruta Chandramouli, Allison Jefferis, Derek A. Applewhite

## Abstract

Allelic variations of Sperm antigen with calponin homology and Coiled-Coil domains 1 Like (SPECC1L) have been associated with a spectrum of cranial-facial pathologies including Teebi hypertelorism and Opitz G/BBB syndrome which manifest as clefting of the palate, wide-eyes, and incomplete closure of the esophagus among others. These pathologies may be indicative of improper cranial neural crest cell (CNCC) delamination and migration. SPECC1L is hypothesized to be an actin-microtubule cross-linking protein as it co-localizes with both microtubules and actin in tissue culture cells. Further, it has been shown to immunoprecipitate with a protein phosphatase complex-1β (PP1β) member, MYPT1, as well as being involved in the PI3K-AKT signaling axis. In this study we sought to investigate the SPECC1L *Drosophila* homolog Spdi and despite sharing close homology with its mammalian counterpart we found that Spdi is associated with both non-muscle myosin-II and actin. RNAi depletion of Spdi led to an increase in focal adhesion dynamics and when we introduced conserved point mutations to Spdi that are analogous to those associated with human disease we observed a further increase in focal adhesion dynamics above that of depletion alone. Collectively, our findings suggest that Spdi is a non-muscle myosin II (NMII) binding protein that likely affects focal adhesion dynamics through this association. Our results also suggest that some of the pathologies associated with allelic variants of SPECC1L may be the result of aberrant cell-matrix adhesion.

## INTRODUCTION

Cellular migration during embryogenesis is a tightly regulated and dynamic process critical for proper development. Among the key players in this process are cranial neural crest cells (CNCCs). CNCCs, which are found at the border of the neural plate in developing animal embryos, emerge around the time of neural tube closure (Martik and Bronner, 2017). These multipotent cells proliferate, migrate ventrally, and ultimately fuse with the core mesoderm and epithelial cover to form the facial primordia—precursors to the facial skeleton and muscles (Jiang et al., 2009; (Jiang *et al*., 2009; Piacentino *et al*., 2020). Understanding the molecular mechanisms underlying CNCC migration is essential for elucidating the etiology of various congenital syndromes that result from defects in this process.

During neurulation, CNCCs undergo an epithelial-to-mesenchymal transition (EMT), transforming from a stationary epithelial sheet into migratory mesenchymal cells (Ferrer-Vaquer *et al*., 2010; Piacentino *et al*., 2020). This transition is orchestrated by cytoskeletal elements such as actin and non-muscle myosin II (NMII), which facilitate the delamination of CNCCs from the dorsal neural tube, enabling their subsequent migration (Martik and Bronner, 2017). Once released from the neural tube, CNCC migration is driven by various signals and pathways, including contact inhibition of locomotion (CIL) mediated by Rho and Rac signaling, chemotaxis through C3a and C3aR signaling, and mechanical stimuli from the underlying mesoderm that modulate cadherin expression (Carmona-Fontaine *et al*., 2008; Theveneau *et al*., 2010; Scarpa *et al*., 2015; Roycroft *et al*., 2018; Shellard *et al*., 2018; Shellard and Mayor, 2019). The mesoderm below CNCCs in a developing embryo stiffens which provides a mechanical stimulus received by an integrin/vinculin/talin complex in CNCCs which results in greater expression of N-Cadherin and a lower expression of E-Cadherin (Scarpa *et al*., 2015).

As cells reach where the face will develop, fusion between the medial nasal, lateral nasal, and maxillary processes occurs. Defects in this fusion can cause a number of craniofacial pathologies, most notably cleft lip with or without cleft palate (Jiang *et al*., 2009), which are associated with a wide range of mutations. A critical protein implicated in CNCC delamination and migration is Sperm antigen with calponin homology and Coiled-Coil domains 1 Like (SPECC1L)(Bhoj *et al*., 2015; Saadi *et al*., 2023). SPECC1L which associates with both actin filaments and microtubules through its calponin homology (CH) domain and coiled-coil domain 2 (CCD2) throughout the cell cycle (Mehta *et al*., 2023; Saadi *et al*., 2023). SPECC1L deficiency has been associated with the increased stability of adherens junctions and reduced rates of CNCC delamination, highlighting its crucial role in this developmental process (Zhang et al., 2020). In congruence with its crucial role in CNCC migration, mutations in SPECC1L have been associated with a spectrum of congenital craniofacial disorders, including Teebi hypertelorism and Opitz G/BBB syndrome (Bhoj *et al*., 2015; Saadi *et al*., 2023).

To further investigate the functional dynamics of SPECC1L, we turned to its *Drosophila melanogaster* homolog, Split Discs (Spdi)(Saadi *et al*., 2011). Initial studies demonstrated that RNAi depletion of Spdi, resulted in impaired cellular adhesion and migration phenotypes, such as crumpled wings and cleft probosci (Walsh and Brown, 1998; Brown *et al*., 2000), mirroring the effects of SPECC1L mutations in humans (Saadi *et al*., 2011). Genetic and phenotypic comparisons between SPECC1L and Spdi reveal a high degree of sequence similarity, particularly in the residues surrounding the putative actin-binding CH domain (Saadi *et al*., 2011; Zhang *et al*., 2020).

Our study aims to characterize the role of Spdi in cytoskeletal regulation. Preliminary data indicate a previously unreported colocalization of Spdi with the heavy chain of NMII, suggesting a tighter connection to myosin than previously proposed. Moreover, mutations in Spdi which are homologous to those in SPECC1L linked to familial forms of craniofacial pathologies resulted in more pronounced effects on focal adhesion turnover than RNAi-mediated depletion alone, implying a possible dominant-negative mechanism (Mehta *et al*., 2023). By elucidating these functional roles and interactions of Spdi and their impact on CNCC migration, we aim to enhance our understanding of the molecular underpinnings of congenital craniofacial disorders and the broader principles of cytoskeletal regulation during embryogenesis.

## RESULTS

### The Drosophila SPECC1L homolog Split discs associates with actin and non-muscle myosin II

A bioinformatic query of the *Drosophila* genome CH domain-containing proteins led to several hits including CG13366, also known as Split Discs (Spdi) (Saadi *et al*., 2011). Spdi is predicted to have three coiled-coil domains (CC1-CC3) separated by linker regions with a CH domain at its C-terminus (Figure 1A) (Saadi *et al*., 2011; Zhang *et al*., 2020). Spdi shares approximately 30% homology with mammalian SPECC1L based on Clustal protein alignments (Madeira *et al*., 2024). The initial characterization of SPECC1L found that EGFP-tagged versions of the protein co-localize with acetylated microtubules while an antibody generated against human SPECC1L co-localized with F-actin suggesting that protein may function as an actin-microtubule cross-linker (Saadi *et al*., 2011). We generated N-terminally fluorescently-tagged versions of Spdi and expressed them off of stable plasmids in *Drosophila* S2R+ cells (Figure 1B-D). We observed a localization pattern more consistent with that of NMII and actin and did not observe microtubule localization as previously reported (Figure 1B & C). To further characterize this expression pattern we expressed fluorescently-tagged Spdi in three additional *Drosophila* cells lines, ML-DmBG3-c2 (DBG3) which are derived from the central nervous system, ML-Dm25-c2 (D25) cells which are derived from third instar wing discs (Ui *et al*., 1987), and Ras^V12^;*wts*^RNAi^ (Ras^V12^) cells which were generated from whole embryo homogenates (Simcox *et al*., 2008). In all cell lines, we observed a localization pattern similar to that of NMII and actin (Figure 1E-G). Thus, it appears that Spdi is an actin-NMII binding protein rather than an actin-microtubule crosslinking protein in *Drosophila* cells.

**Figure 1.**
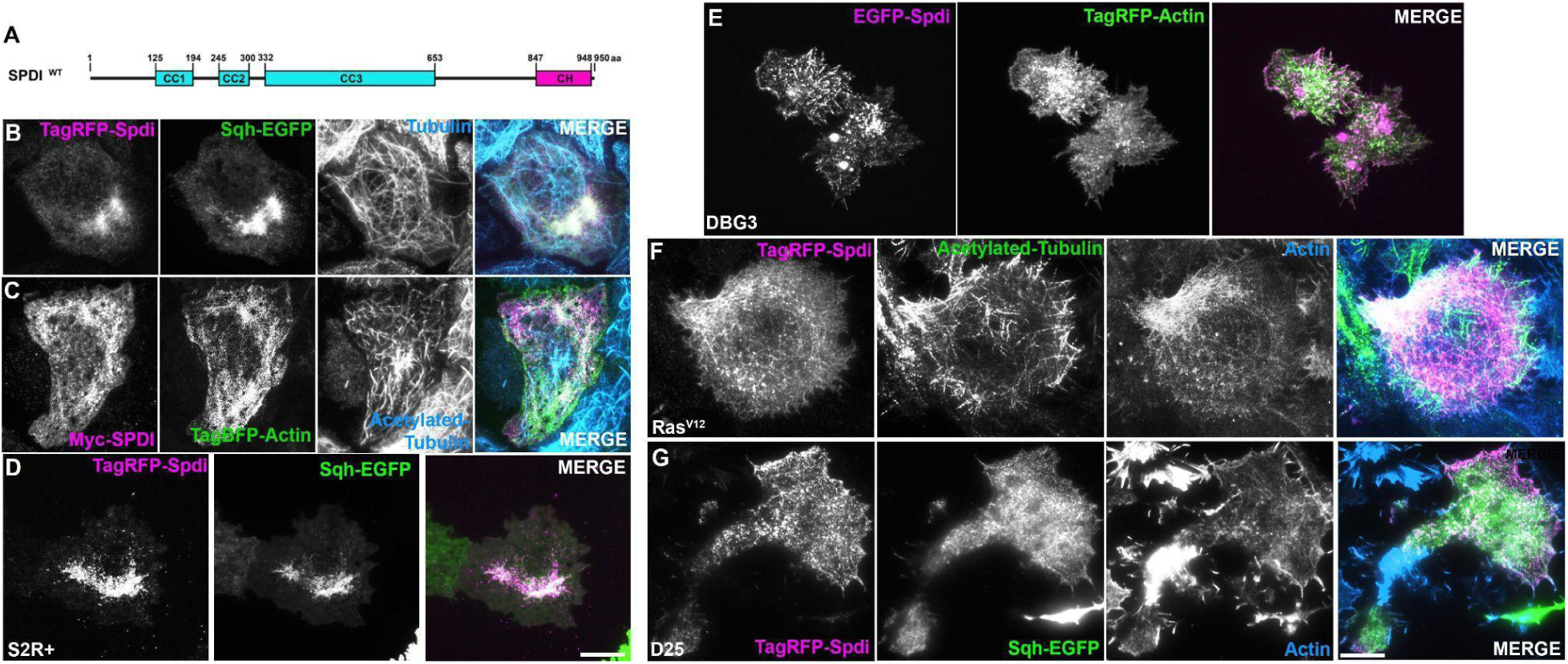
The *Drosophila* SPECC1L homolog, Split discs appears to associate with non-muscle myosin II and actin and not tubulin. (A) Domain organization of Split discs (Spdi). Shown in cyan are Spdi’s three coiled-coil domains (CC1-CC3), and in magenta is Spdi’s calponin homology (CH) domain. Numbers indicate amino acid positions from N-terminus to C-terminus. (B-C) *Drosophila* S2R+ cells imaged by total internal reflection (TIRF) microscopy, scale bar 10 µm. (B) S2R+ cell transfected with TagRFP-Spdi (magenta in merged image) and Spaghetti squash (Sqh)-EGFP (green in merged image) fixed and immunostained for beta-Tubulin (cyan in merged image). (C) S2R+ cell transfected with myc-tagged Spdi (magenta in merged image) and TagBFP-actin (green in merged image) fixed and immunostained for acetylated Tubulin (cyan in merged image). (D) Live-cell imaging of an S2R+ cell transfected with TagRFP-Spdi (magenta in merged image) and Sqh-EGFP (green in merged image). (E) A *Drosophila* DBG3 cell derived from the central nervous system, co-expressing EGFP-Spdi (magenta in merged image) and TagRFP-actin (green in merged image). (F) A *Drosophila* Ras^V12^ cell derived from homogenized embryos and are epithelial in nature transfected with TagRFP-Spdi (magenta in merged image) and fixed and immunostained for acetylated Tubulin (green in merged image) and phalloidin to mark actin (cyan in merged image). (G) A *Drosophila* D25 cell derived from the third instar wing disc transfected with TagRFP-Spdi (magenta in merged image) and Sqh-EGFP (green in merged image) fixed and stained with phalloidin to mark actin (cyan in merged image). Scale bar 10 µm.

The NMII holoenzyme is composed of two heavy chains (known as Zipper in flies), two essential light chains (Mlc-c), and two regulatory light chains (Spaghetti Squash or Sqh) and in an effort to better identify which subunit of NMII Spdi potentially associates with we depleted each of these polypeptides in *Drosophila* S2R+ cells and co-expressed TagRFP-Spdi with EGFP-Sqh (Figure 2A-C) or EGFP-Zipper (Figure 2F). Depletion of Zipper predictably led to a dispersion and disruption of the peri-nuclear NMII network (Figure 2A & B), however it also appears to shift Spdi to what is likely actin as it was observed to more heavily decorate bundles in the interior of the cell a well as a population of putative actin filaments in the transition zone of these cells (Figure 2B). Similarly, depletion of Mlc-c and Sqh led to a disruption of the NMII network however unlike Zipper depletion, Spdi remained associated with the dispersed NMII puncta (Figure 2C & D). We depleted myosin heavy chain-like (Mhc1), also known as Myosin-18 which co-polymerizes with NMII in mammalian systems (Billington *et al*., 2015) (Figure 2E). Depletion of Mhc1 did not appear any different than that of control RNAi treated cells (Figure 2A) suggesting that if Spdi is associating with NMII it is not through Myosin-18. Depletion of the protein phosphatase complex-1 (PP1) targeting subunit, myosin binding subunit (Mbs), results in hyper-phosphorylated NMII pool (Vasquez *et al*., 2014). This leads to a less dispersed and more pronounced perinuclear NMII network. Spdi localization also shifts with NMII upon MBS depletion (Figure 2F). Collectively, these results suggest that Spdi associates with NMII and actin rather than being an actin-microtubule crosslinker as what is suggested for its mammalian homolog. Further this association is likely through NMII heavy chain given the shift in localization from NMII to actin following Zipper depletion.

**Figure 2.**
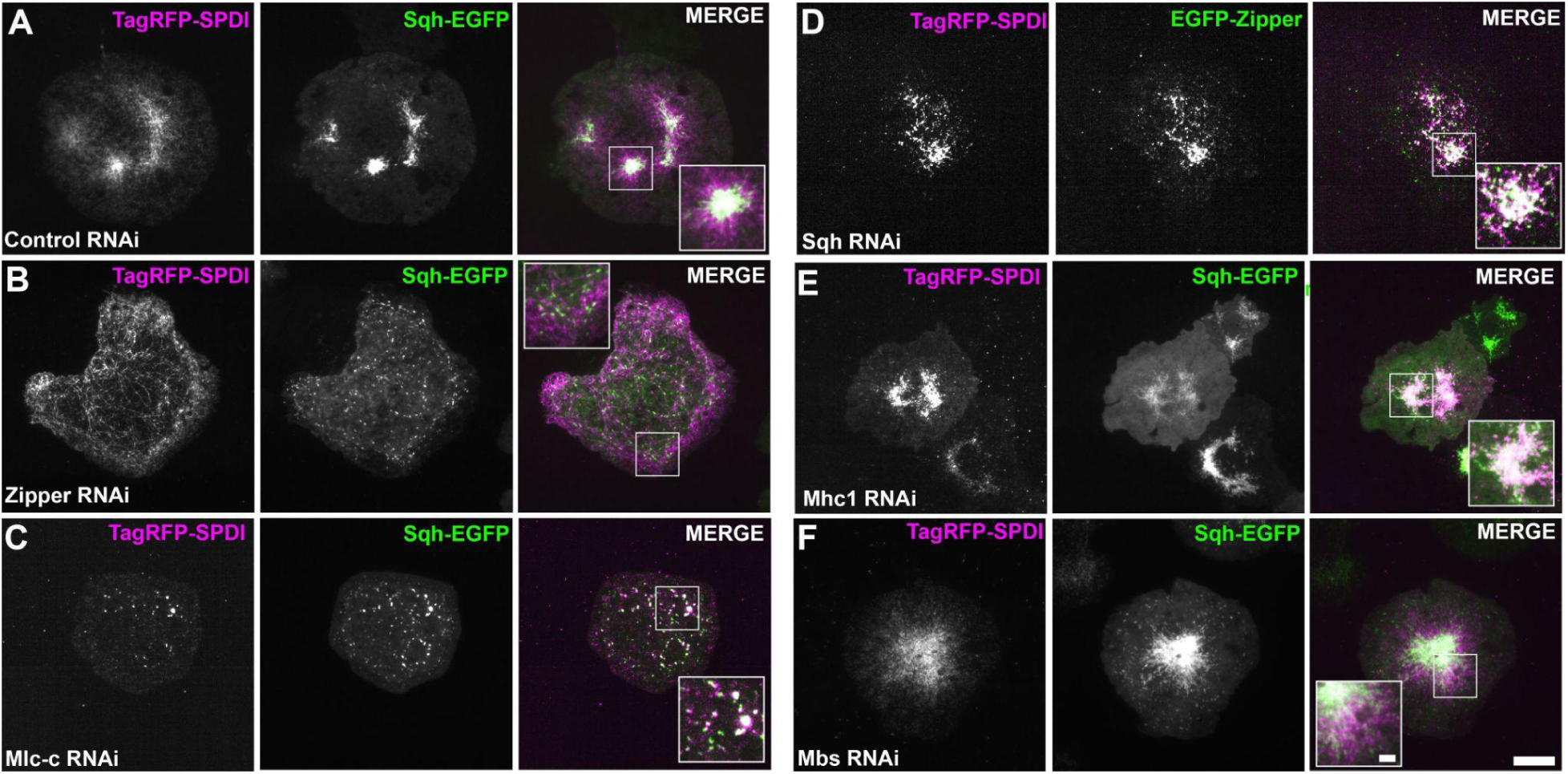
RNAi depletion suggests that Spdi associates with NMII heavy chain Zipper. (A-C) Live-cell imaging by TIRF microscopy of *Drosophila* S2R+ cells transfected with TagRFP-Spdi (magenta in merged image) and Sqh-EGFP following treatment with RNAi targeting (A) control, (B) Zipper, and (C) Myosin light chain-cytoplasmic (Mlc-c, essential light chain). (D) A *Drosophila* S2R+ cell transfected with TagRFP-Spdi (magenta in merged image) and EGFP-Zipper (green in merged image) following RNAi depletion of Sqh. (E & F) *Drosophila* S2R+ cells transfected with TagRFP-Spdi and Sqh-EGFP following RNAi treatment with (E) Myosin heavy chain-like 1 (Mhc1) and, (F) the PP1 complex member Myosin binding subunit (Mbs). Scale bar 10 µm.

### Spdi’s CH domain is required but is not sufficient for NMII localization

We next took a structure-function approach in order to identify which domains of Spdi are required for its localization to NMII. Spdi is composed of three coiled-coil (CC1-3) domains and a C-terminal calponin homology (CH) domain (Figure 3A). We generated a series of TagRFP-tagged deletion constructs, SpdiΔCCD1 (a deletion of residues 1 to 194), SpdiΔCCD1,2 (a deletion residues 1 to 300), SpdiΔCH (a deletion of residues 847 to 950), and SpdiCHD (a construct containing residues 847 to 950) and co-expressed them with EGFP-tagged Sqh (Figure 3A-F). Prior to co-transfection, *Drosophila* S2R+ cells were treated with RNAi targeting the 3’-untranslated region (3’UTR) of Spdi for seven days in order to deplete the endogenous pool of Spdi. The efficacy of our RNAi was quantified using RT-qPCR with depletion using Spdi RNAi (coding sequence) and 3’UTR Spdi resulting in 81.97% and 70.74% knockdown, respectively (Supplemental Figure 1A). Currently there are no antibodies against Spdi, so to confirm these RT-qPCR results, we generated a stable S2R+ cell line that expresses a Myc-tagged Spdi under the control of a metallothionein promoter. We then treated these cells with Spdi or Spdi 3’UTR RNAi and then induced expression of Myc-tagged Spdi and performed a Western blot on whole cell lysates (Supplemental Figure 1B). As expected, treatment with Spdi RNAi led to depletion of the Myc-tagged Spdi while RNAi targeting the 3’UTR Spdi was unable to target our exogenously expressed fusion protein.

**Figure 3.**
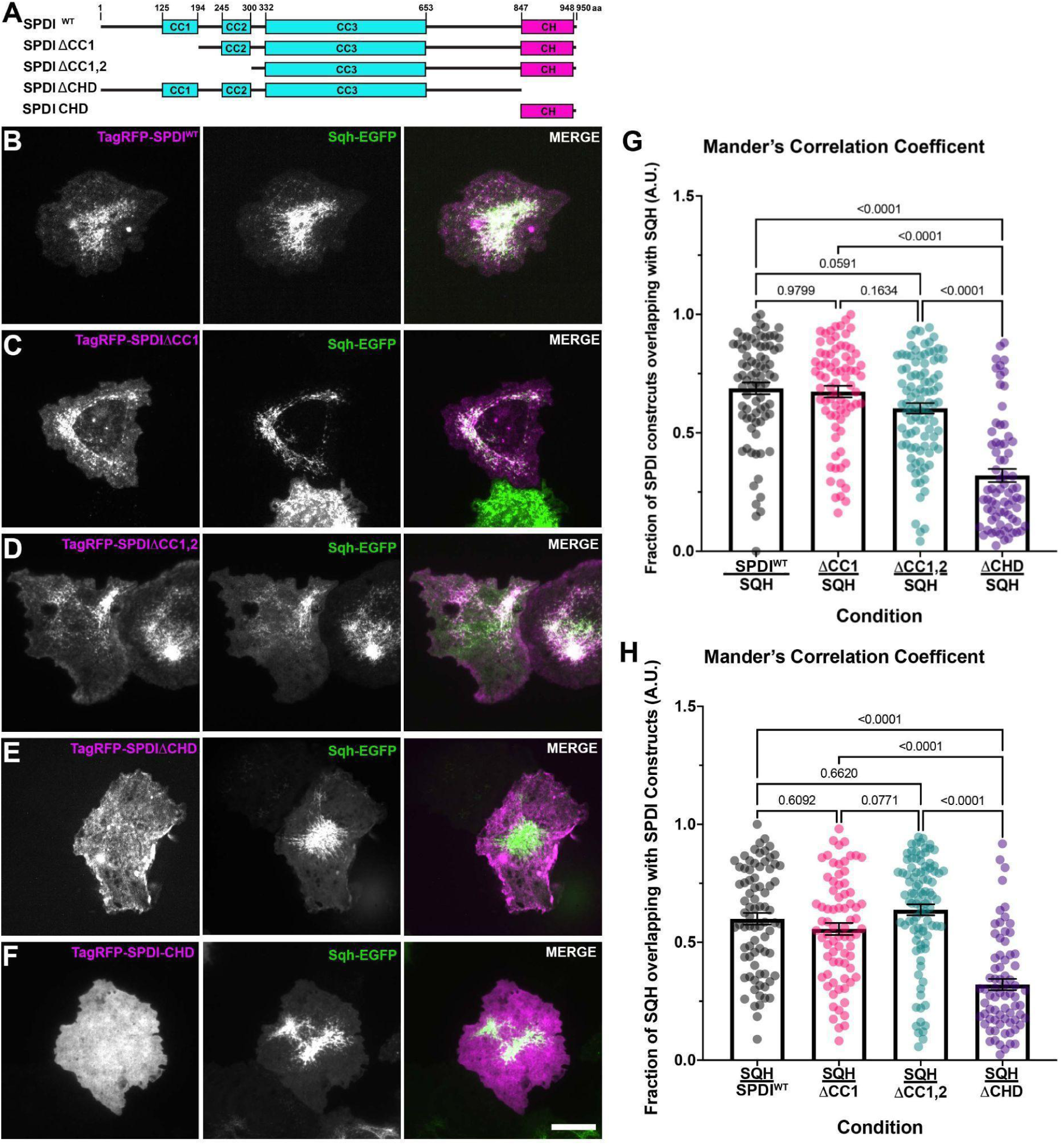
Spdi’s C-terminal CHD is needed for colocalization with NMII. (A) The domain organization of the deletion constructs used to identify the requirement for NMII colocalization. The various deletion constructs are as follows: SpdiΔCCD1; deletion of residues 1-194, SpdiΔCCD1,2; deletion residues 1-300, SpdiΔCHD; deletion of residues 847-950, and SpdiCHD; residues 847-950 only. (B-F) Live-cell imaging of *Drosophila* S2R+ cells imaged by TIRF microscopy following a seven-day course of Spdi 3’UTR RNAi treatment and co-transfection of Sqh-EGFP (green in merged image) and (B) full-length TagRFP-Spdi (magenta in merged image), (C) TagRFP-SpdiΔCCD1(magenta in merged image), (D) TagRFP-SpdiΔCCD1,2 (magenta in merged image), (E) TagRFP-SpdiΔCHD (magenta in merged image), and (F) TagRFP-SpdiCHD (magenta in merged image). Scale bar 10 µm. (G & H) Bar and scatter plots of Mander’s Correlation Coefficient for (G) the fraction of Spdi overlapping with Sqh, and (H) the fraction of Sqh overlapping with Spdi. Full-length Spdi is shown in gray circles, SpdiΔCCD1 is shown in pink circles, SpdiΔCCD1,2 shown in cyan circles, and Spdi-CHD is shown in purple circles. There was a statistically significant difference in the means of Spdi-CHD and Sqh as compared to the other constructs (*p*-values are shown on the graphs, One-way ANOVA with Tukey’s post-hoc analysis, N = 3, n = 72-95 cells).

Using Mander’s correlation coefficient we then quantified the degree of colocalization between these deletion constructs and NMII (Figure 3G & H). In comparing the amount of overlap between Spdi and Sqh or the inverse (the amount of Spdi overlapping with Sqh) deletion of CC1 alone or in combination with CC2 was not statistically significantly different from the amount of overlap between full-length wild-type Spdi and Sqh (Figure 3G & H). However, expression of SPDIΔCH did show a statistically significant reduction in the amount of overlap between Spdi and Sqh as compared to both full-length Spdi, SPDIΔCC1, and SPDIΔCC1,2. It did appear to associate with actin rich portions of the cell rather than being completely cytoplasmic (Figure 3E). Thus, it appears that the CH domain is the determining factor in the colocalization between Spdi and NMII and as such we sought to test whether expression of Spdi CH domain was sufficient to target NMII. CH domains are traditionally categorized as type 1, 2, or 3 with type 1 having the intrinsic ability to bind actin, type 2 usually working in tandem with type 1 to form a CH1-CH2 high affinity actin binding domain, and type 3 diverging the most, capable of binding actin but also able to interact with a number of other proteins (Stradal *et al*., 1998; Yin *et al*., 2020). We aligned Spdi’s CH domain alongside SPECC1L’s CH domain and compared them to archetypal CH1, CH2, and CH3 domains from human alpha-actinin, MICAL, and Calponin respectively. In comparing the amino acid sequence of Spdi’s CH domain, it appears to more closely align with that of CH2 domains, having the shared SSSW sequence (Supplemental Figure 1C). We expressed a TagRFP-tagged Spdi CH domain and compared its localization to that of Sqh as a marker of NMII and observed a largely cytoplasmic localization for this construct similar to that of expression of an untagged EGFP molecule. This result suggests that Spdi may oligomerize in order to increase its affinity for the cytoskeleton, rely on another cytoskeletal binding protein for its localization, or has a cryptic domain that functions alongside this CH2 domain to facilitate cytoskeletal association.

### Spdi has a slight preference for NMII in the “open” phosphorylated state

NMII undergoes cycles of activation where its regulatory light chain is phosphorylated at residues threonine 20 and threonine 21 resulting in an open, oligomerization competent state, and cycles of inactivation where it is dephosphorylated resulting in a closed state where it is unable to oligomerize and interact with actin. We wondered if Spdi has a preference for either of these states and to test this we again turned to Mander’s correlation coefficient to quantify the degree of colocalization. We co-expressed TagRFP-Spdi with either wild-type EGFP-Sqh, a non-phosphorylatable regulatory light chain mutant, EGFPSqhAA (T20A,T21A), or a phosphomimetic mutant, EGFP-SqhEE (T20E, T21E) and compared the degree of colocalization to that of cells co-transfected with EGFP-Sqh and TagRFP-Sqh as a control (Supplemental Figure 2A-D). We observed a statistically significant increase in the amount of colocalization between Spdi and SqhAA and SqhEE as compared to wild-type Sqh both when comparing the overlap between Spdi and Sqh and the inverse (Supplemental Figure 2E & D). While it is surprising that both of these conditions lead to a slight increase, it should also be noted that these experiments were performed in the presence of endogenous Sqh, thus we are observing the localization of both labeled and unlabeled NMII oligomers. However, there was a slight, but statistically significant difference between the fraction of Spdi overlapping with SqhEE as compared to Spdi overlapping with SqhAA suggesting that Spdi may favor the phosphomimetic version of Sqh (Supplemental Figure 2E).

### Spdi likely associates with NMII through its coiled-coil tail domain

Our RNAi depletion experiments (Figure 2C & D) suggest that Spdi’s NMII association is not dependent on regulatory light chain (Sqh) nor the essential light chain (Mlc-c) as their depletion did not lead to loss of colocalization with NMII, whereas depletion of Zipper led to a distinct shift in localization (Figure 2B). This likely indicates that Spdi associates with Zipper either directly or indirectly rather than the other polypeptides of the holoenzyme. Using an allelic series of Zipper deletion constructs, we co-expressed TagRFP-Spdi with EGFP-tagged full length Zipper, Zipper Neck-Rod (residues 775-1972), Zipper RodΔNterm (residues 909-1972), and Zipper HMM-EGFP (residues 1-910), and again used Mander’s correlation coefficient to quantify the degree of colocalization (Figure 4A) (Franke *et al*., 2005). In quantifying the amount of overlap between Zipper and Spdi we observed a drastic, statistically significant decrease in the amount of colocalization in between Spdi and Zip HMM as compared to all other conditions (Figure 4F). Conversely, the degree of colocalization between Spdi and the allelic Zipper series (the amount of Spdi overlapping with Zipper) was statistically indistinguishable for wild-type Zipper, Zip Neck-Rod, Zipper RodΔNterm (Figure 4G). There was a slight but statistically significant increase in the colocalization of Zip HMM as compared to Zip Neck-Rod, but upon visual inspection, the Zip HMM construct appears to be distributed all over the cell in a nonspecific manner similar to what has been observed when free EGFP is expressed (Figure 4E). These results taken together with our RNAi depletion experiments indicate that Spdi likely associates with NMII through its coiled-coil/ tail domain.

**Figure 4.**
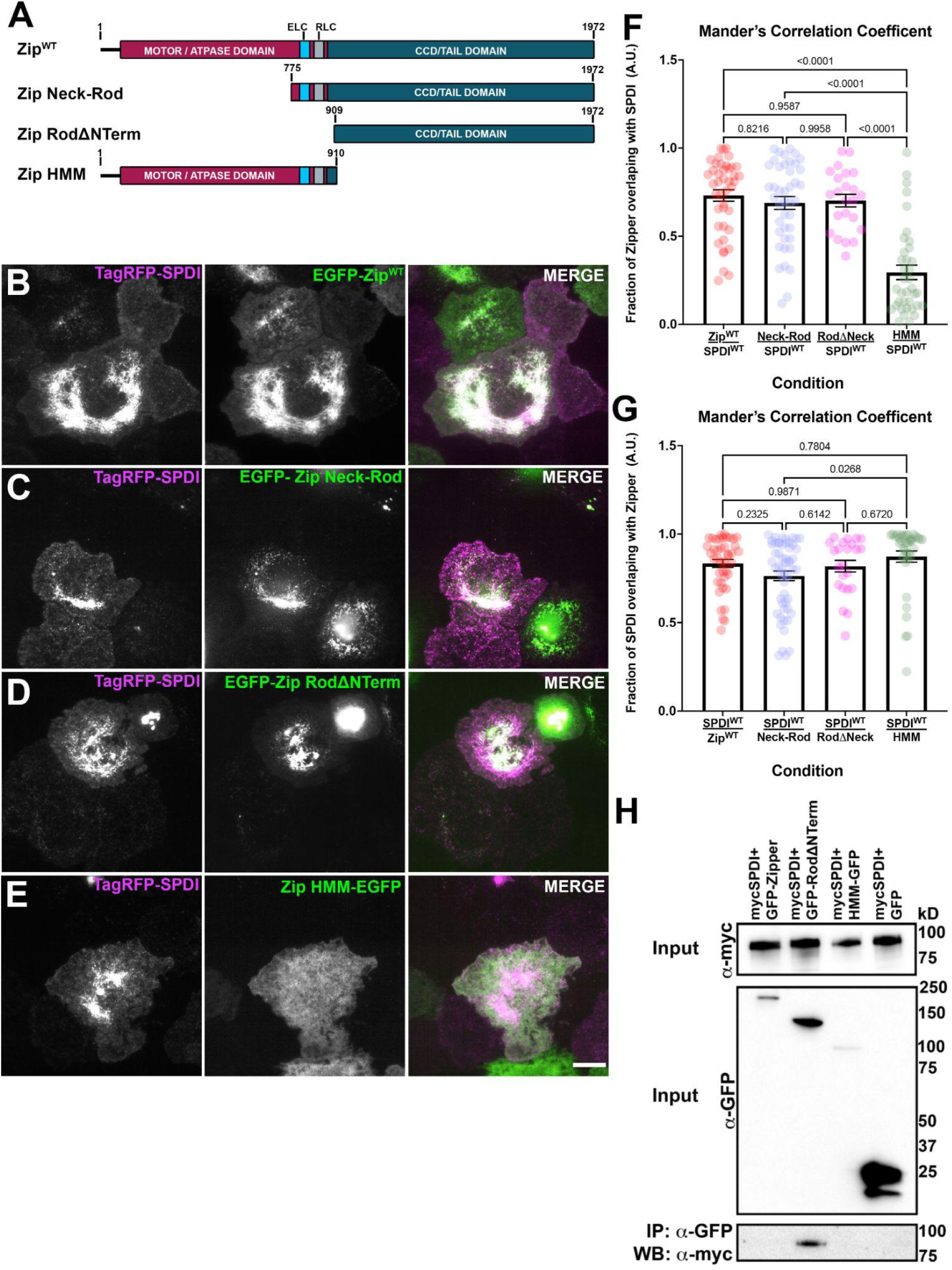
Spdi preferentially associates with the coiled-coil tail domain of NMII. (A) Domain organization of *Drosophila* Zipper and an allelic series used to identify the requirement for Spdi colocalization. Maroon, the motor/ATPase domain, and dark cyan, the coiled-coil/tail domain. The essential light chain binding site is in cyan, and the regulatory light chain binding site is shown in gray. The allelic series is as follows: Zipper Neck-Rod (residues 775-1972), Zipper RodΔNterm (residues 909-1972), and Zipper HMM (residues 1-910). (B-E) Live-cell imaging by TIRF microscopy of *Drosophila* S2R+ cells co-transfected with TagRFP-Spdi (magenta in merged image) and (B) EGFP-Zipper (full length, green in merged image), (C) EGFP-Zipper Neck-Rod (green in merged image), (D) EGFP-Zipper RodΔNterm (green in merged image), and (E) Zipper HMM-EGFP (green in merged image). Scale bar 10µm. (F & G) Scatter and bar plots of Mander’s Correlation Coefficients for the amount of overlap between (F) Zipper constructs and Spdi, and (G) Spdi and the Zipper constructs. Red circles represent full length Zipper and Spdi, light blue circles, Zipper Neck Rod and Spdi, pink circles, Zipper RodΔNterm and Spdi, and the moss circles, Zipper HMM and Spdi. There was a statistically significant decrease in the amount of overlap between Zipper HMM and Spdi (*p-*values are shown on graphs, One-way ANOVA with Tukey’s post-hoc analysis, N = 3, n = 23-54 cells). (H) Immunoprecipitation of *Drosophila* S2R+ whole cell lysates co-transfected with myc-Spdi and EGFP-Zipper, EGFP-Zipper RodΔNterm, EGFP-Zipper HMM, and EGFP only as a non-specific control. We observed and interaction between EGFP-Zipper RodΔNterm and myc-Spdi and no other conditions.

To further test if Spdi is associating with the NMII rod/tail domain, we performed an immunoprecipitation (Figure 4H). We coexpressed a myc-tagged Spdi with either EGFP-tagged full length Zipper, Zipper RodΔNterm, Zip HMM, or a construct expressing untagged EGFP and pulled down with an anti-GFP antibody. We then performed a Western blot with an anti-myc antibody. EGFP-Zipper RodΔNterm was able to pull down myc-Spdi while full length Zipper, Zipper HMM-EGFP, and GFP-only failed to pull down myc-Zipper. The Zipper RodΔNterm construct lacks the binding sites for both the ECL and the RLC, further suggesting that Spdi associates with the NMII coiled-coil/tail domain. While the full length Zipper construct also contains the coiled-coil/tail domain, the paracrystals/aggregates that can form when the heavy chain and regulatory light chain are out stoichiometry as a result of overexpression may preclude the binding of other proteins (Franke *et al*., 2006). In support of our immunoprecipitation results, Goering and colleagues found that wild-type mouse SPECC1L from embryonic tissues was able to co-immunoprecipitate with NMIIA and NMIIB (Goering *et al*., 2021). Although mammalian SPECC1L does not colocalize with NMII in cells, both Spdi and SPECC1L may each form a complex with NMII.

### A Glutamine-to-Proline mutation at residue 266 decreases Spdi’s association with NMII

Allele variations of SPECC1L are associated with a spectrum of cranial-facial disorders which are putatively linked to aberrant cranial neural crest cell migration and adhesion (Bhoj *et al*., 2015; Kruszka *et al*., 2015; Wilson *et al*., 2016). We generated three Spdi point mutants, glutamine 266 to proline (Q266P), glycine 915 to serine (G915S), and arginine 931 and glutamine (R931Q), which are analogous SPECC1L alleles associated with cranial-facial pathologies in humans (Saadi *et al*., 2011; Bhoj *et al*., 2015; Wilson *et al*., 2016) (Figure 5A). To test if these point mutants co-localize with NMII we first treated S2R+ cells with dsRNA targeting the 3’-untranslated region (3’UTR) of Spdi to deplete the endogenous pool. We then co-expressed TagRFP-tagged Spdi mutants with EGFP-tagged Zipper and used Mander’s coefficient to assess the degree of colocalization (Figure 5B-G). Both the Q266P mutation and the R931Q mutation lead to a statistically significant decrease in the fraction of overlap observed between Spdi and Zipper and the overlap observed between Zipper and Spdi (Figure 5F & G), however this decrease was far more pronounced in cells expressing the Q266P mutation (Figure 5C). The amount of colocation between Zipper and the G915S mutant was no different from that of wild-type or our TagRFP-EGFP Sqh control (Figure 5B, D, F & G). In addition to investigating how these Spdi point mutants colocalize with wild-type NMII, we wondered if they may confer differences in associating with NMII in the “closed” or “opened” confirmation. We treated S2R+ cells with 3’UTR Spdi dsRNA and then co-expressed TagRFP-tagged Spdi Q266P, G915S, or R931Q with either SqhWT-EGFP, SqhAA-EGFP, or SqhEE-EGFP and determined the degree of colocalization via Mander’s coefficient (Supplemental Figure 3A-L). We observed a co-localization pattern similar to wild-type NMII and these SPDI mutants, in that regardless the state of the regulatory light chain, both the Q266P and the R931Q mutants show a decrease in the amount of colocalization with NMII, while G915S mutant co-localization was more similar to that of wild-type SPDI, albeit to a somewhat lesser degree (Supplementary Figure 3M & N). These differences were statistically significant (Supplemental Table 1). These results suggest that the disease associated alleles colocalize NMII in the “open” and “closed” conformations in the same manner as wild-type Spdi.

**Figure 5.**
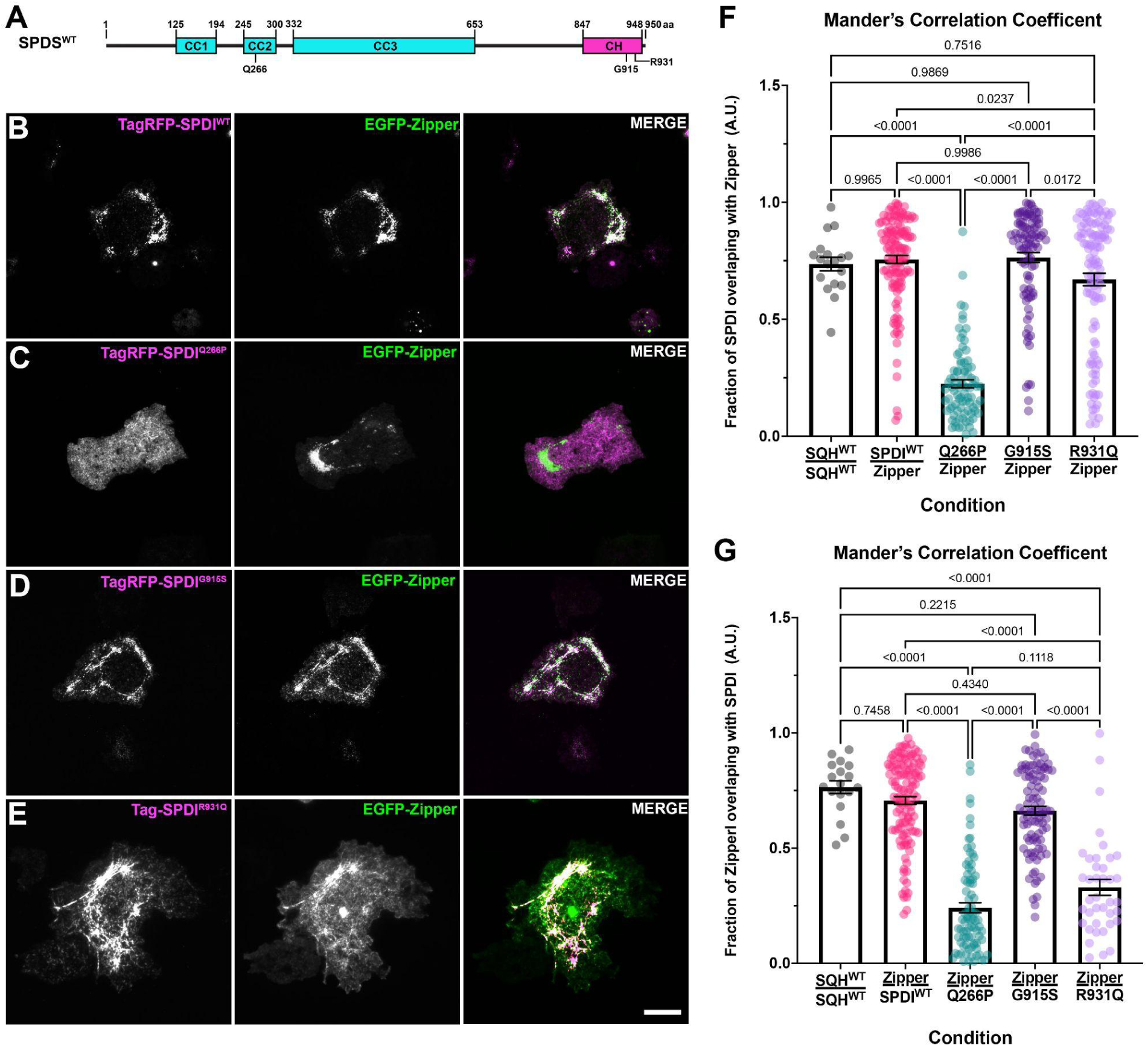
Disease associated allelic variant of Spdi decreases the degree of colocalization with NMII. (A) Spdi domain organization indicating the positions of the conserved point mutations associated with cranial-facial pathologies. (B-E) Live-cell imaging by TIRF microscopy of *Drosophila* S2R+ cells co-transfected with EGFP-Zipper (green in merged images) and (B) TagRFP-Spdi^WT^ (wild-type Spdi, magenta in merged image), (D) TagRFP-Spdi^Q266P^(Spdi Q266P, magenta in merged image),(E) TagRFP-Spdi^G915S^ (Spdi G915S magenta in merged image), and TagRFP-Spdi^R931Q^ (Spdi R931Q magenta in merged image). Scale bar 10 µm. (F & G) Scatter and bar plot quantifying Mander’s Correlation Coefficient for the degree of overlap between (F) Spdi and Zipper and (G) Zipper and Spdi. As a positive control we included the degree of overlap between Sqh-EGFP and Sqh-TagRFP (gray circles). The quantification in overlap between wild-type Spdi and Zipper is shown in pink circles, Spdi Q266P in cyan circles, Spdi G915S and Zipper in purple circles, and Spdi R931Q and Zipper in lavender circles. There was a statistically significant decrease in the degree of overlap between Zipper and Spdi Q266P and Zipper and Spdi R931Q (*p*-values are shown on graphs, One-way ANOVA with Tukey’s post-hoc analysis, N = 3, n = 18-119 cells).

### Depletion of Split discs increases focal adhesion dynamics

The cranial-facial pathologies in humans associated with atypical alleles of SPECC1L suggest that it may play a role in regulating CNCC cell migration and adhesion (Saadi *et al*., 2011; Wilson *et al*., 2016). In further support of this, escapers of amorphic alleles of Spdi in *Drosophila* also displayed phenotypes that are often associated with aberrant cell migration and adhesion (Prout *et al*., 1997; Brown *et al*., 2000; Franke *et al*., 2010). In particular, the wing blistering phenotype observed in some escapers phenocopies mutations in focal adhesion proteins in this tissue. These results motivated us to examine focal adhesion dynamics in migratory *Drosophila* tissue culture cells. We transfected *Drosophila* DBG3 cells with mCherry-Vinculin as a marker for focal adhesions and tracked their assembly and disassembly by TIRF microscopy (Figure 6A & B). It should be noted that modEncode data suggests that Spdi is expressed relatively high in DBG3 cells (Landt *et al*., 2012). Depletion of Spdi in DBG3 cells led to a statistically significant increase in rate constants for both focal adhesion assembly and disassembly (Figure 6C & D). Similarly, we tracked focal adhesion dynamics in D25 cells which are epithelial in nature. In these cells we observed a statistically significant increase in the focal adhesion disassembly rate constant while focal adhesion assembly rate constant was no different from that of control cells (Figure 7A & B). We also measured the rates of random cell migration for both cell types. Depletion of Spdi in DBG3 cells resulted in a slight but statistically significant increase in the speed of random cell migration while depletion of Spdi in D25 cells resulted in a statistically significant slower migration speed (Figure 7C & D). It should be noted that cell migration speeds in D25 cells following Spdi knock-down were bi-modal in nature, with approximately 40% of the cells migrating faster than the mean with the remaining 60% at or below the mean. Further, D25 cells migrate one order of magnitude faster than DBG3 cells (Figure 7C & D). This heterogeneity in focal adhesion dynamics and random cell migration speeds can be expected given the distinct tissues from which these cells are derived and the differing rates of random cell migration. Collectively, these results suggest that Spdi may play a role in regulating cell-matrix adhesion dynamics.

**Figure 6.**
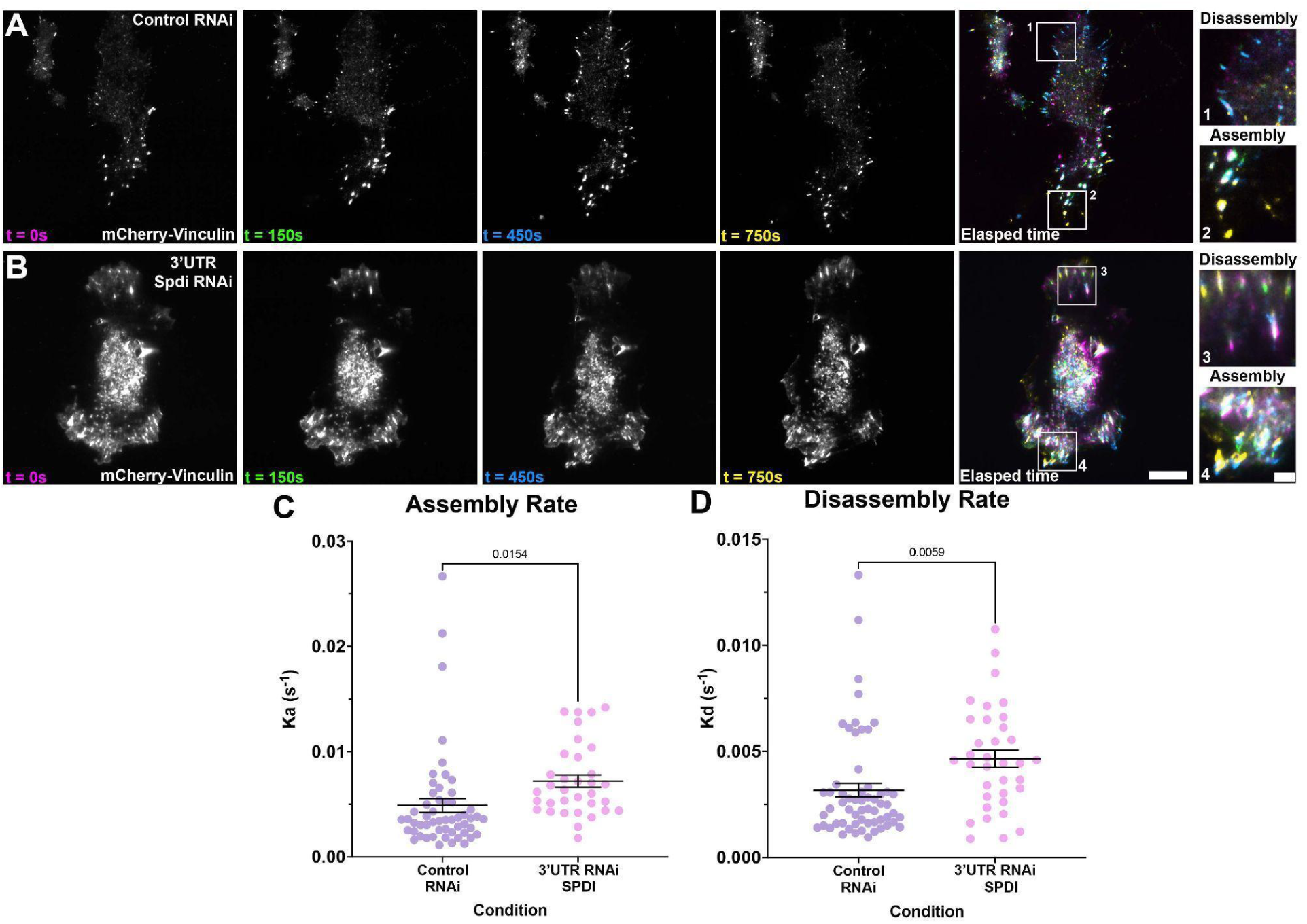
Depletion of Spdi leads to increased focal adhesion dynamics. (A & B) Live-cell imaging by TIRF microscopy of *Drosophila* DBG3 cells expressing mCherry-Vinculin following (A) control RNAi or (B) 3’UTR Spdi RNAi. Time points are as follows: 0 seconds (magenta in merged elapsed time image), 150 seconds (green in merged elapsed time image), 450 seconds (blue in merged elapsed time image), and 750 seconds (yellow in merged elapsed time image). White boxes (1-4) in merged elapsed time images indicate (1 & 3) focal adhesion assembly and (2 & 4) focal adhesion disassembly and are shown at higher magnification. Scale bars are 10 µm in lower magnification images and 2 µm in high magnification images. (C & D) Scatter plots of (C) focal adhesion assembly rates and (D) disassembly rates following control RNAi (lavender circles) and 3’UTR Spdi RNAi (pink circles). There was a statistically significant increase in both focal adhesion assembly and disassembly as the results of Spdi depletion (*p*-values are shown on graphs, Student’s t-test, N = 3, n = 34-54 focal adhesions [assembly] and 35-59 focal adhesions [disassembly]).

**Figure 7.**
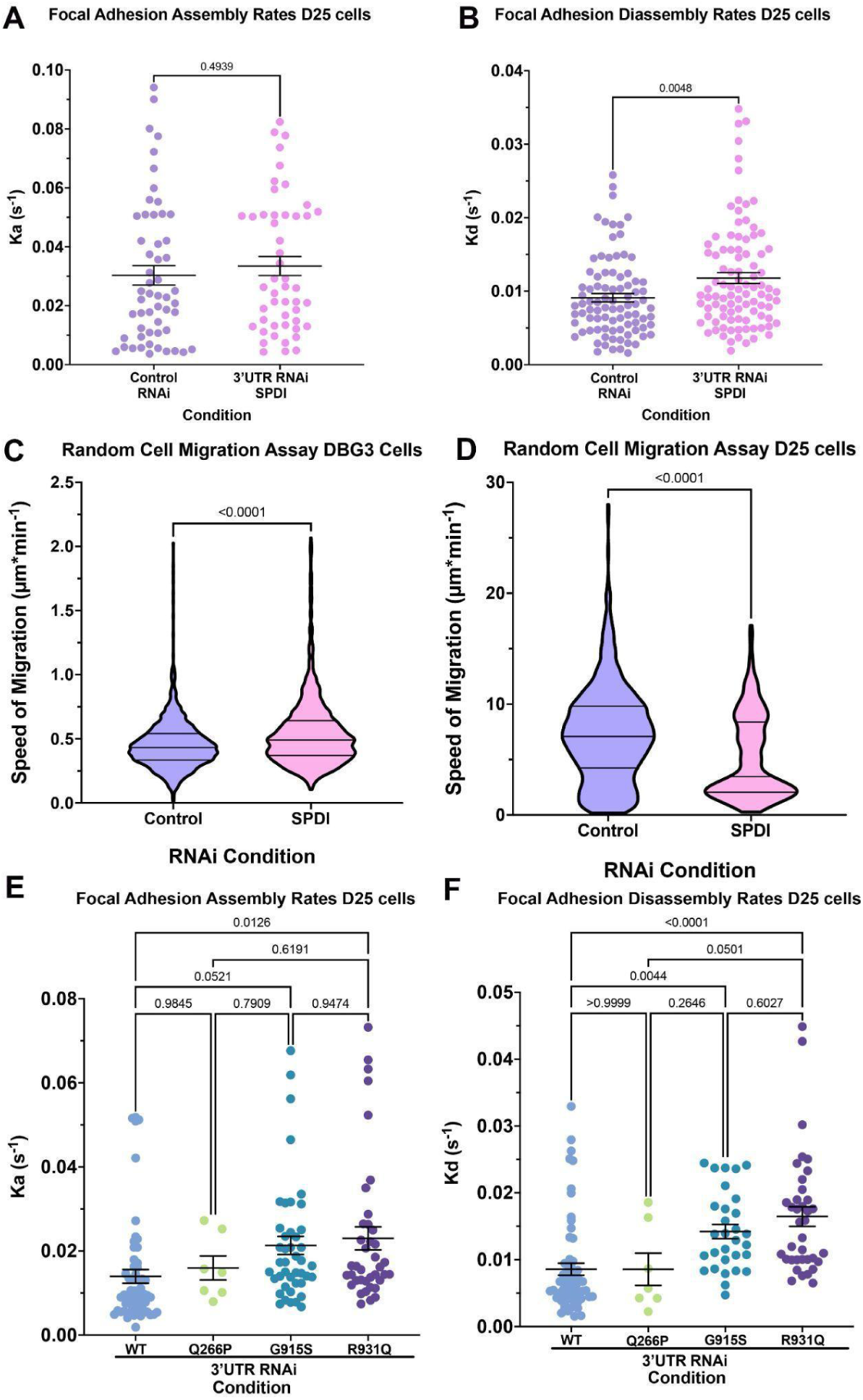
Focal adhesion and cell migration rates are altered by depletion of Spdi or expression of disease associated alleles. (A & B) Scatter plots of focal adhesion (A) assembly rates and (B) disassembly rates in D25 cells following treatment with control RNAi (purple circles) or 3’UTR Spdi RNAi (pink circles). Focal adhesion assembly rates were no different between control and 3’UTR Spdi RNAi treated cell, however, focal adhesion disassembly rates were statistically significantly increased following depletion of Spdi (*p*-values shown on graphs, Student’s t-test, N = 3, n = 88-95 focal adhesions). (C & D) Violin plots of the rates of random cell migration for (C) DBG3 cells and (D) D25 cells following treatment with control (purple) or 3’UTR Spdi (pink) RNAi. There was a statistically significant increase in the rate of DBG3 random cell migration (*p*-value shown on graph, Student’s t-test, N = 3, n = 1113-1456 cells), while Spdi depletion lead to a statistically significant decrease in D25 random cell migration (*p*-value shown on graph, Student’s t-test, N = 3, n = 625-629 cells). (E & F) Focal adhesion (E) assembly and (F) disassembly rates following treatment with 3’UTR Spdi RNAi and re-expression of wild-type Spdi (light blue circles), Spdi Q266P (lime green circles), Spdi G915S (cyan circles), and Spdi R931Q (purple circles). Expression of Spdi R931Q led to a statistically significant increase in focal adhesion assembly (*p*-value shown on graph, One-way ANOVA with Tukey’s post-hoc analysis, N = 3, n = 7-58 focal adhesions). Expression either Spdi G915S or Spdi R931Q led to a statistically significant increase in focal adhesion disassembly (*p*-values shown on graphs, One-way ANOVA with Tukey’s post-hoc analysis, N = 3, n = 7-37 focal adhesions).

While depletion of Spdi appears to increase focal adhesion dynamics, the cranial-facial pathologies observed in some families are associated with atypical SPECC1L alleles and not the loss of gene expression. Using our Spdi constructs that contain these mutations to conserved residues (Figure 5A), we depleted DBG3 cells of endogenous Spdi using 3’UTR Spdi RNAi and then added back either wild type EGFP-tagged Spdi, or EGFP-Spdi Q266P, G915S, and R931Q alongside mCherry-Vinculin to track focal adhesion dynamics (Figure 8A-D). While there was no statistically significant increase in focal adhesion assembly between wild-type and the Q266P allele, we did observe a statistically significant increase in focal adhesion assembly rate constants in the G915S and R931Q point mutants. Further the focal adhesion assembly in the R931Q allele was statistically faster than focal adhesion assembly rates in cells expressing the G951S allele (Figure 8E). Similarly, focal adhesion disassembly was also altered by expression of these disease associated alleles. We observed a statistically significant increase in focal adhesion disassembly as compared to wild-type in DBG3 cells expressing all three disease associated alleles (Figure 8F). In this case, there was a stepwise increase in disassembly rates, increasing in the Q266P allele and continuing in the G915S and R931Q alleles. Again, we performed parallel experiments using D25 cells, and observed similar trends. Expression of both G915S and R931Q alleles following treatment with 3’UTR Spdi RNAi lead to a statistically significant increase in focal adhesion dynamics as compared to cells expressing wild-type or the Q266P allele variant (Figure 7E & F). These disease associated alleles potentially alter cell-matrix adhesion dynamics beyond that of Spdi depletion alone and may indicate a dominant negative mechanism of action.

**Figure 8.**
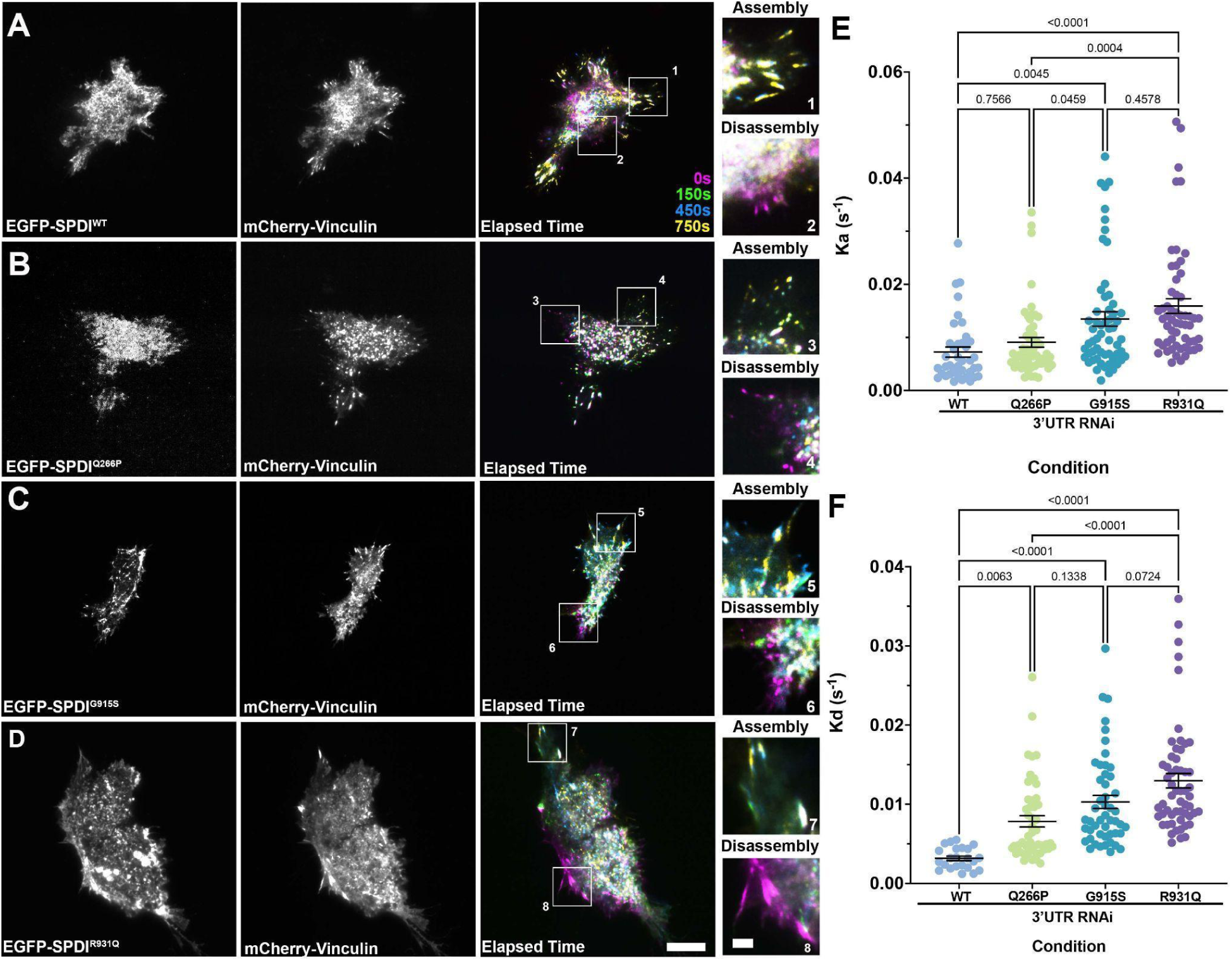
Expression of conserved disease associated alleles leads to altered focal adhesion dynamics. (A-D) Live-cell TIRF imaging of *Drosophila* DBG3 cells, co-expressing mCherry Vinculin, and (A) wild-type Spdi, (B) Spdi Q266P, (C) Spdi G915S, or (D) Spdi R931Q. Elapsed time merged images indicate time points as follows: 0 seconds (magenta in merged elapsed time image), 150 seconds (green in merged elapsed time image), 450 seconds (blue in merged elapsed time image), and 750 seconds (yellow in merged elapsed time image). White boxes in merged images (1-8) indicate regions of focal adhesion (1,3,5,7) assembly and (2,4,6,8,) disassembly shown at higher magnification. Scale bars 10 µm in low magnification images and 2 µm in high magnification images. (E and F) Scatter plots of (E) focal adhesion assembly rates and (F) focal adhesion disassembly rates following treatment with 3’UTR Spdi RNAi and expression of wild-type Spdi (light blue circles), Spdi Q266P (lime green circles), Spdi G915S (turquoise circles) and Spdi R931Q (purple circles). Upon depletion and endogenous Spdi and re-expression of Spdi G915S and R931Q we observed a statistically significant increase in focal adhesion dynamics (*p*-values are shown on graphs, One-way ANOVA, N = 3, n = 39-58 focal adhesions [assembly] and n = 49-56 [disassembly]).

## DISCUSSION

Familial patterns of cranial-facial pathologies led to the identification of SPECC1L where point mutations to this protein correlated to what is now described as a spectrum of cranial-facial disorders (Saadi *et al*., 2011; Kruszka *et al*., 2015; Bhoj *et al*., 2019; Zhang *et al*., 2020). Further interrogation of SPECC1L homologs in mice, zebrafish, *Drosophila*, and human tissue culture cells further defined a role for this protein as an actin-microtubule crosslinking protein that plays putative roles in regulating adhesion, localization of the myosin phosphatase complex, and the PI3K-AKT signaling axis (Wilson *et al*., 2016; Hall *et al*., 2020; Goering *et al*., 2021; Mehta *et al*., 2023; Saadi *et al*., 2023). The initial studies identified SPECC1L as an actin-microtubule cross-linking protein that strongly associates with acetylated tubulin leading microtubule stabilization. Deletion of its C-terminal CH domain prevented this actin-microtubule cross-linking-dependent stabilization and reduced its association with microtubules (Saadi *et al*., 2011). Similarly, SPECC1L allelic variations associated with craniofacial pathologies had reduced microtubule localization as did a SPECC1L CRISPR mutant carrying a deletion in the second coiled-coil domain. This deletion in SPECC1L shifted the protein’s localization to perinuclear actin bundles (Hall *et al*., 2020; Goering *et al*., 2021). This is particularly curious as the second coiled-coil domain appears to be a hotspot for craniofacial pathologies. SPECC1L also localizes to cell-cell junctions in U2OS and its knock-down leads to increased actin stress fibers and a disruption of adherens junction markers E-cadherin, myosin IIB, and beta-catenin. Further, these adherens junction markers were disrupted *in vivo* in mouse knock-outs, which was concomitant with a failure of neural tube closure and the delamination of cranial neural crest cells (Wilson *et al*., 2016). This increase in actin stress fiber may be indicative of an overall increase in cellular contractility. In support of this hypothesis, a series of mass spec and immunoprecipitation experiments found that SPECC1L directly interacts with protein phosphatase complex-1β (PP1β) member MYPT1 in U2OS cells (Mehta *et al*., 2023). This interaction was dependent on the third coiled-coil domain of SPECC1L and the C-terminal coiled-coil and leucine zipper domains of MYPT1 and it is thought that SPECC1L dynamically regulates the localization of the MYPT1/PP1β complex between actin and microtubules (Mehta *et al*., 2023). Thus, SPECC1L potentially brings with it a phosphatase complex that can counter NMII contractility. Finally, both pan-AKT, and active phospho-AKT immunostaining were reduced in SPECC1L deficient tissues *in vivo* (Wilson *et al*., 2016). Given the potential antagonistic relationship between AKT and E-cadherin, this finding further implicates SPECC1L in the regulation of cell-cell adhesion (Larue and Bellacosa, 2005).

In our investigation of the *Drosophila* SPECC1L homolog Spdi, we find that it robustly colocalizes with NMII and actin rather than actin and microtubules (Figures 1 & 2). This interaction is dependent on the rod/coiled-coil domain of the heavy chain of NMII and deletions of the first and second coiled-coil domains of Spdi did little to change this localization. It was only upon deletion of the C-terminal CH domain of Spdi did we observe a significant decrease in NMII co-localization (Figures 3 & 4). While this localization differs somewhat from that of its mammalian counterpart, several observations support an interaction between SPECClL and NMII. Firstly, both wild-type SPECC1L and a CH domain deletion variant co-immunoprecipitated with both NMIIA and NMIIB (Goering *et al*., 2021), and as discussed above, SPECC1L also co-immunoprecipitated with MYPT1 a phosphatase that is hypothesized to regulate NMII contractility (Mehta *et al*., 2023). Given the degree of conservation between SPECC1L and Spdi we were able to introduce point mutations analogous to those allelic variations of SPECC1L associated with cranial-facial pathologies. Parallel to what has been observed in mouse and human tissue culture cells between SPECC1L and microtubules (Saadi *et al*., 2011), we found that the Q266P point mutant (which corresponds to Q415P in human SPECC1L) greatly reduced the localization of Spdi with NMII (Figure 5). In addition, we found a slight but statistically significant decrease in the colocalization between the R931Q (which corresponds to R1098Q) point mutant and NMII (Figure 5). Residue Q266/Q415 is in the second coiled-coil domain and when CRISPR was used to generate in-frame deletions of CCD2 in mice, heterozygotes behaved in a way the was characteristic with an autosomal dominant with increases in the occurrence of ventral body wall closure defects, cleft palate, and exencephaly (Goering *et al*., 2021). This in-frame CCD2 deletion also showed increased localization to actin bundles and a decrease in localization to microtubules which may be more reminiscent of actin/NMII localization (Goering *et al*., 2021).

Another parallel that can be drawn between mammalian SPECC1L and *Drosophila* Spdi is their potential roles in regulating adhesion. In the case of mouse SPECC1L, deficient tissue resulted in increased stabilization of junction markers. This in turn could lead to the defects in delamination and neural tube closure observed in knockout mice (Wilson *et al*., 2016). Our study implicated Spdi in the regulation of focal adhesion dynamics with RNAi depletion leading to a mild increase in both focal adhesion assembly and disassembly (Figure 6). However, when we tracked focal adhesions in cells expressing point mutations associated with cranial-facial pathologies, we observed a more robust stepwise increase in both focal adhesion assembly and disassembly starting with a point mutant in the second CCD (Q266P) and ending with a point mutant in the C-terminal CH domain (R931Q) (Figure 8). Focal adhesion maturation is highly dependent on NMII contractility, so the increase in focal dynamics could be attributed to Spdi’s putative regulation of NMII contractility (Giannone *et al*., 2007; Burnette *et al*., 2011; Elosegui-Artola *et al*., 2016; Yolland *et al*., 2019). Cell migration is dependent on optimal adhesion so changes in focal adhesion dynamics lead to changes in the rates of cell migration (Gupton and Waterman-Storer, 2006). We previously found that depletion of the actin-microtubule crosslinking protein Short stop (Shot) led to an increase in focal adhesion dynamics which then corresponded with an increase in cell migration (Zhao *et al*., 2022). Upon Spdi depletion we observed both an increase in cell trajectory speed in DBG3 cells which are derived from the central nervous system, and a decrease in cell speed in D25 cells which are derived from third instar wing discs (Figure 7). While these results are contradictory from what we observed following Shot depletion and internally between these two cell lines, other factors should be considered. Firstly, D25 and DBG3 cells are from different tissues and have very different rates of migration with D25 cells being about one order of magnitude faster. Secondly, there was a bimodal distribution of migration rates in D25 cells with a large proportion of cells migrating much faster than control depleted cells which is more in line with what we observed upon depletion of Shot. Finally, Shot depletion was concomitant with an increase in lamellipodial dynamics, we did not measure lamellipodial dynamics in Spdi depleted cells. We did however observe a similar stepwise increase in focal adhesion dynamics upon expression of Spdi point mutants associated with cranial-facial pathologies across both cell lines despite having different tissues of origin (Figures 7 & 8).

## Conclusion

We found that the *Drosophila* homolog of SPECC1L, Split Discs, localizes to NMII and actin. This localization is different from what has been observed for SPECC1L in mammalian systems as it was found to be an actin-microtubule cross-linking protein. Further, we find that depletion of Split Discs leads to increases in focal adhesion assembly and disassembly, but moreover, expression of point mutants associated with cranial-facial pathologies led to an increase in focal adhesion dynamics above that of depletion suggesting a possible dominant negative mechanism. We speculate that Split Discs’ role in focal adhesion dynamics is a result of its regulation of NMII contractility.

## Materials and Methods

### Plasmid constructs

*split discs* (*spdi*) and or the *myc* tag was cloned into pMT/HisA (Thermo Fisher Scientific, Waltham, MA) by traditional cloning methods to generate pMTSPDSHisA and pMTMycSPDS. Inserts *egfp* and *tagRFP* were cloned into pMTSPDSHisA by traditional cloning techniques. Gene inserts were amplified by cloning primers by KOD polymerase. The reactions were pooled and precipitated by isopropanol and sodium acetate. For the generation of *birA* and fluorescent gene-tagged constructs, all PCR products and vectors were digested (by manufacturer’s recommended instructions) in CutSmart buffer and KpnI-HF (New England Biolabs, Ipswich, MA), with the addition of CIP (Thermo Fisher Scientific) in the vector reactions. The digested products were gel purified by Zymoclean Gel DNA Recovery Kit (Zymo Research, Tustin, CA) by the manufacturer’s instructions. Inserts were ligated to the vectors using Rapid DNA Ligation Kit (Sigma, St. Louis MO) by the manufacturer’s instructions, except incubated at 22°C for 8 min. NEB5α competent *E. coli* (High-Efficiency) cells (New England Biolabs) were transformed with the ligation reactions by the manufacturer’s instructions. Plasmids were checked for accuracy by sequencing (ACGT, Inc., Wheeling, IL).

### Cell Culturing

For detailed cell culture instructions see (Rogers et al., Applewhite et al). In brief, *Drosophila* ML-DmDBG3-c2, ML-DmD25-c2, and Ras^V12^;*wt^s^*^RNAi^ (DBG3, D25, and Ras^V12^ cells) were obtained from Drosophila Genomics Resource Center, Bloomington, IN) and maintained a 25°C in Schneider’s Drosophila Medium (Gibco #21720, Waltham, MA) supplemented with 10 µg/ml insulin (Gibco #12585-014), 10% FBS (Thermo Fisher Scientific), and 1X Antibiotic-Antimycotic (Gibco #15240-062). Cells were grown in cell-culture flasks and passed every 4–5 days. *Drosophila* S2R+ cells were cultured at 25 °C in Shields and Sang media (S&S; Sigma S8398, 0.5 g/l NaHCO3, 1 g/l yeast extract, 2.5 g/l bactopeptone) with 10% fetal bovine serum (FBS) (Thermo Fisher Scientific) and 1 × Antibiotic–Antimycotic (Thermo Fisher Scientific; 15,240–062. Cells were grown in cell-culture flasks and passed every 4–5 days. Transfections were carried out using FuGENE HD Transfection Reagent (Promega, Madison, WI) and was used according to the manufacturer’s instructions. Briefly, FuGENE reagent (6 µl) was added to 2-4 µg plasmid DNA in 100 µl water and mixed by immediately pipetting. Solutions were allowed to incubate at room temperature for no more than 5 minutes, then was added dropwise to each 9.6 cm^2^ well, along with 0.6-1 mM CuSO_4_ (if induction was required) and incubated at 25°C for 1-2 days prior to imaging. For RNAi Treatments cells were cultured to 50-90% confluency in a 9.6 cm^2^ well in 1 ml of media. To this 1 ml of cell and media, 1 µl (approximately 100 ng of dsRNA) was added daily for 5-7 days.

### Immunofluorescence and imaging

For S2R+ cells, concanavalin A (con A, Sigma) was applied to glass bottom dishes (1.5 glass coverslips attached to laser-cut 35-mm tissue culture dishes with UV-curable adhesive (Norland Products, Cranbury, NJ), removed, and the dishes were left to air dry. Once dry, cells in Shang’s and Shields media were allowed to attach for 45 min. For DBG3, D25, and Ras^V12^ cells were plated on glass-bottom dishes treated with ECM harvested from the cells as described in (Currie and Rogers, 2011) in modified Schneider’s Drosophila Medium (above) and allowed to attach for at least 60 min. For immunofluorescence, cell were fixed with a 10% paraformaldehyde solution (Electron Microscopy Sciences, Hatfield, PA) in PEM buffer, 100 mM PIPES, 1 M MgCl2, 1 mM EGTA, pH to 6.8) for 15 min at room temperature. The cells were rinsed 3 × with Phosphate-buffered Solution (PBS; 20 mM Tris, 150 mM NaCl, pH 7.4), and cells were blocked with 5% goat serum in PBS for 10 min. The block was removed and a 1:200 dilution rabbit anti-acetylated tubulin primary antibody (1:200) in 5% goat serum in PBS was added and left to incubate at room temperature for one hour, or at 4 °C overnight. The cells were rinsed 3 × with PBS and then incubated with a solution of 1:100 anti-Rabbit Alexa Fluor 598 secondary antibody (Jackson ImmunoResearch Laboratories, Inc., West Grove, PA), with 1:100 Alexa Fluor 488 Phalloidin and incubated in the dark at room temperature for one hour. The cells were rinsed 3 × with PBS, and mounted using Dako Fluorescence Mounting Medium (Agilent, Santa Clara, California). Cells were imaged using a Nikon Eclipse Ti-E inverted microscope (Nikon, Tokyo, Japan) using a 100x/1.49NA oil immersion TIRF objective.

### Quantification of focal adhesion assembly and disassembly rates

DBG3 and D25 cells were transfected with mCherry-Vinculin (Ribeiro *et al*., 2014) under the control of the metallothionein promoter, which served as a marker for focal adhesions, and then imaged every 5 s over a 15-min period. The image analysis software Imaris (Bitplane, Concord, MA) was used to measure the intensity of the focal adhesions over time. The fluorescence of an area close to the focal adhesion was measured and subtracted from the focal adhesion fluorescence values in order to account for background fluorescence. To smooth out variation between different frames, a running three-frame average was applied to the fluorescence values of each frame (Stehbens *et al*., 2014). These values were then fitted against known equations for assembly and disassembly, with disassembly modeled using single exponential decay and assembly represented as a sigmoid, logistic function (Stehbens *et al*., 2014). The assembly (Ka) and disassembly (Kd) constants for graphs that accurately represented changes in fluorescence values over time were then recorded. To corroborate that the values were not artifacts of background fluorescence, the sizes of the fluorescent objects were recorded and any object that was either smaller or larger than conventionally accepted focal adhesion measurements (0.5–10 μm2) was discarded. Images were captured by TIRF microscopy (described above).

### Colocalization analysis

Colocalization was analyzed by line-scan analysis and Mander’s coefficient analysis. For line-scan analysis, a 10 μm line was drawn from the cell edge inward and fluorescence intensity was measured. These values were normalized and then averaged for all cells within that condition. Mander’s coefficient analysis was performed using the Just Another Colocalization Program (JACoP) plug-in for ImageJ (Bolte and Cordelières, 2006). Briefly, intensity thresholds were manually set for both fluorescence channels and then the fraction of overlap was calculated in each direction. Images were captured by TIRF microscopy (described above).

### Random cell migration

DBG3 and D25 cells were plated at a subconfluent density on ECM-coated glass-bottom dishes and allowed to attach overnight. Cells were imaged every 5 min for 6 h by phase-contrast microscopy using a 40×/0.75NA objective. Individual cells were manually tracked using Manual Tracker (ImageJ). Speed is calculated by the total trajectory speed divided by the total time.

Spdi: Split Discs
CNCC: cranial neural crest cell
PP1: phosphatase complex-1
SPECC1L: Sperm antigen with calponin homology and Coiled-Coil domains 1 Like
CIL: contact inhibition of locomotion
CH: calponin homology
CC: Coiled-coil
Sqh: Spaghetti Squash (NMII regulatory light chain)
Zip: Zipper (NMII heavy chain)
Mhc1: myosin heavy chain-like
RLC: regulatory light chain
ELC: essential light chain
UTR: untranslated region

## Acknowledgements

We acknowledge the Drosophila Genomics Resources Center (DGRC) (National Institutes of Health [NIH] grant 2P40OD010949 to the DGRC) and Dan Kiehart (Duke University) for reagents. In addition we thank members of the Applewhite lab and members of the Reed College Physics department including Jay Ewing, Lucus Illing, Darrell Shroeter, and Alison Crocker for their thoughtful discussions and support during the preparation of this manuscript. Further, we thank Greta Glover and Jeff Brown for their support with equipment and reagents. This work was supported by the NIH (R15GM122019-01 to D.A.A) and the Reed College Department of Biology.

## SUPPLEMENTAL FIGURES

**Supplementary Figure 1.**
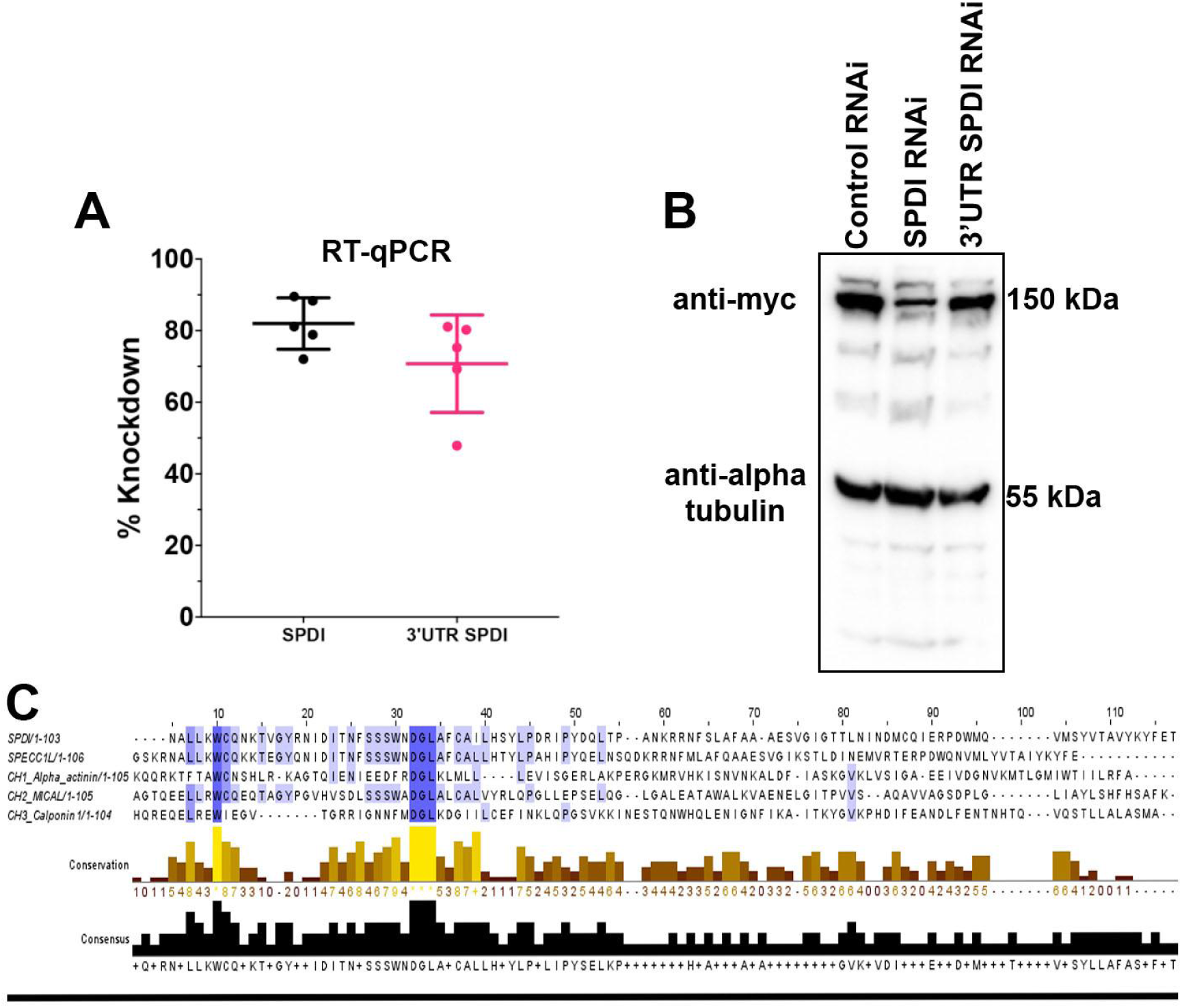
RT-qPCR efficacy and analysis of Spdi’s Calponin homology domain. (A) A scatter plot of RT-qPCR efficacy for Spdi RNAi (coding sequence, black circles) and 3’ untranslated region of Spdi (3’UTR, pink circles) (N = 5). (B) Western blot analysis of whole cell lysates from stable S2R+ cells expression Myc-tagged Spdi under the control of metallothionein promoter (pMT). From left to right, cells treated with control RNAi, Spdi RNAi, and 3’UTR Spdi RNAi. Mouse anti-myc antibody was used to detect Myc-tagged Spdi, and alpha-tubulin was used as a loading control. (C) Alignment of Spdi and SPECC1L CH domains alongside archetypal CH domains, CH1 (Alpha-actinin), CH2 (MICAL), and CH3 (Calponin). The degree of conservation is shown in blue (the darker the blue the more conserved). Conservation is also shown in the histogram below on a 0-9 scale with 0 being the least conserved and 9 being the most (brown to yellow respectively). Asterisks indicate identical residues. The black histogram represents the consensus amino actin sequence.

**Supplemental Figure 2.**
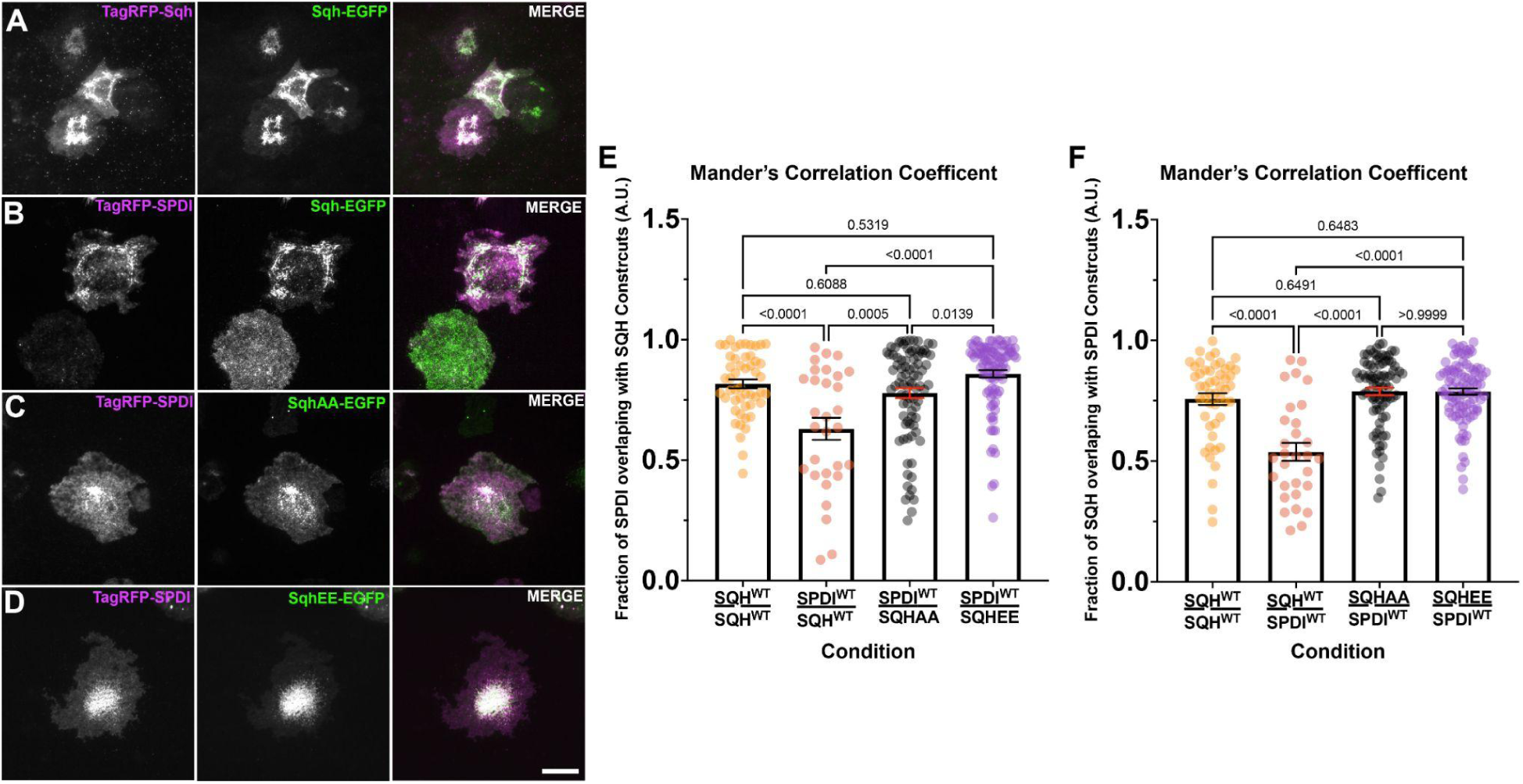
Spdi has a slight preference for phosphomimetic Sqh. (A-D) Live-cell imaging of *Drosophila* S2R+ cells imaged by TIRF microscopy. (A) Co-transfection of TagRFP-Sqh (magenta in the merged image) and Sqh-EGFP (green in the merged image). (B) Co-transfection of TagRFP-Spdi (magenta in the merged image) and Sqh-EGFP (green in merged image). (C) Co-transfection of TagRFP-Spdi (magenta in the merged image) and non-phosphorylatable SqhAA-EGFP (green in merged image). (D) Co-transfection of TagRFP-Spdi (magenta in the merged image) and phosphomimetic SqhEE-EGFP (green in merged image). Scale bar 10 µm. (E & F) Bar and scatter plots of Mander’s Correlation Coefficient quantifying the amount of overlap between (E) Spdi and Sqh (WT, AA, EE) and (F) Sqh (WT, AA, EE) and Spdi. As a positive control we also compared the amount of overlap between wild-type Sqh-TagRFP, and wild-type Sqh-EGFP (orange circles). The overlap between wild-type Spdi and wild-type Sqh is shown in salmon circles, wild-type Spdi and SqhAA in black circles, and wild-type Spdi and SqhEE in purple circles. There was a statistically significant increase in the amount of overlap between Spdi and SqhAA and SqhEE over wild-type Sqh. In addition there was a slight but statistically significant increase in overlap between Spdi and SqhEE over that of SqhAA (*p*-values are shown on graphs, One-way ANOVA with Tukey’s post-hoc analysis, N = 3, n = 30-95 cells).

**Supplementary Figure 3.**
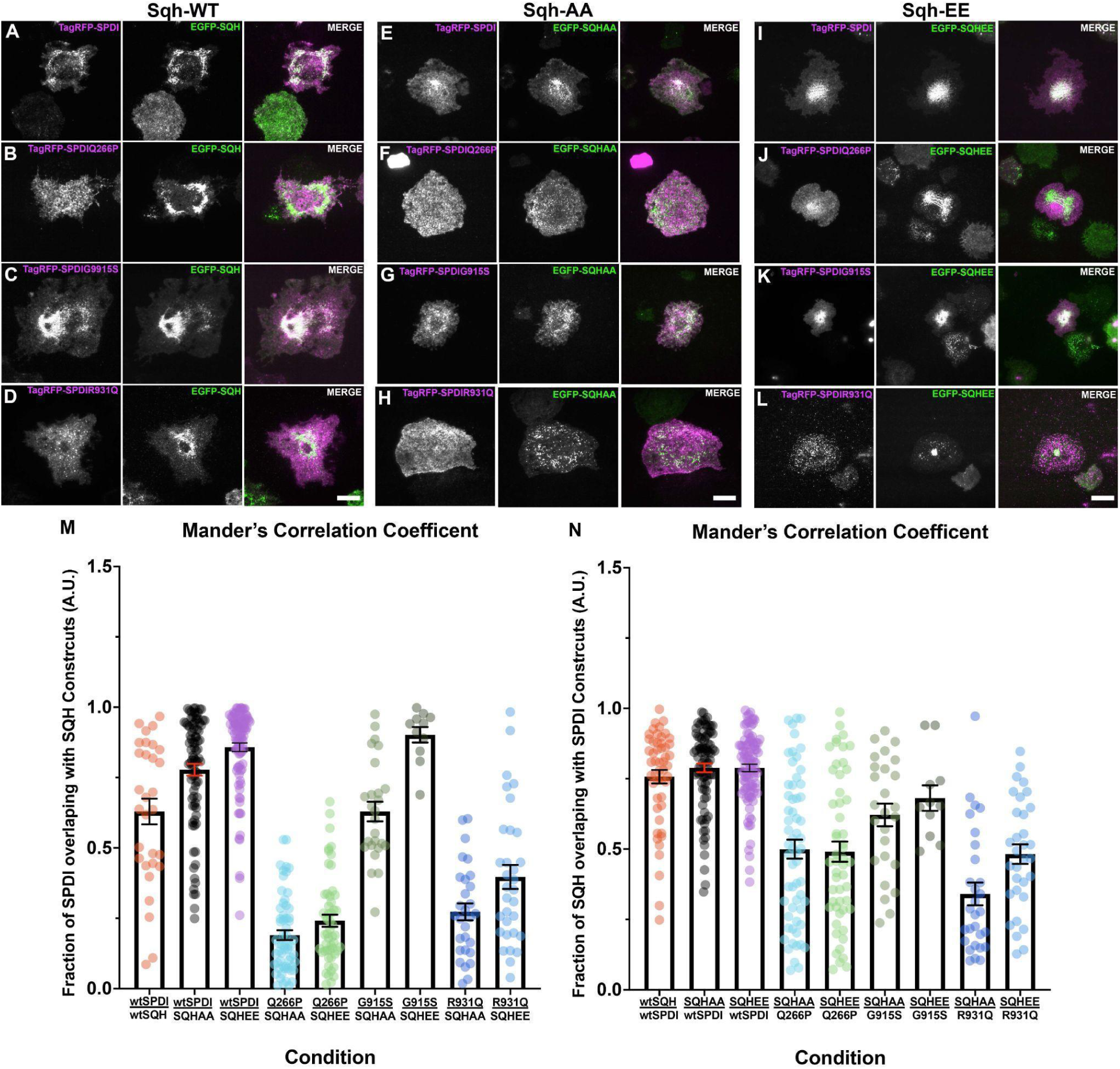
Comparison of Spdi point mutants to the phosphorylation state of the regulatory light chain. (A-L) Live-cell TIRF imaging of *Drosophila* S2R+ cells. (A-D) Co-transfection of Sqh-EGFP (wild-type, green in merged image) with (A) TagRFP-Spdi (wild-type, magenta in merged image), (B) TagRFP-Spdi Q266P (magenta in merged image), (C) TagRFP-Spdi G915S (magenta in merged image), and (D) TagRFP-Spdi R931Q (magenta in merged image). (E-H) Co-transfection of SqhAA-EGFP with (E) TagRFP-Spdi (wild-type, magenta in merged image), (F) TagRFP-Spdi Q266P (magenta in merged image), (G) TagRFP-Spdi G915S (magenta in merged image), and (H) TagRFP-Spdi R931Q (magenta in merged image). (I-L) Co-transfection of SqhEE-EGFP with (I) TagRFP-Spdi (wild-type, magenta in merged image), (J) TagRFP-Spdi Q266P (magenta in merged image), (K) TagRFP-Spdi G915S (magenta in merged image), and (L) TagRFP-Spdi R931Q (magenta in merged image). Scale bars 10 µm. (M & N) Scatter and bar plots of Mander’s Correlation Coefficients quantifying the degree of overlap between (M) Spdi and Sqh constructs, and (N) Sqh constructs and Spdi. Salmon circles is the overlap between Spdi (wild-type) and Sqh (wild type), black circles, Spdi (wild type) and SqhAA, purple circles, Spdi (wild-type) and SqhAA, cyan circles, Spdi Q266P and SqhAA, lime green circles, Spdi Q266P and SqhEE, moss green circles, Spdi G915S and SqhAA, gray circles Spdi G915S and SqhEE, dark blue circles, Spdi R931Q and SphAA, and light blue circles Spdi R931Q and SqhEE.

**Supplemental Table 1.** Statistical Analysis^a^ of the comparison between Spdi’s point mutants and the regulatory light chain’s phosphorylation state.

| Spdi/Sqh<sup>b</sup> | *p*-value | Spdi/Sqh<sup>b</sup> | *p*-value | Sqh/Spdi<sup>c</sup> | *p*-value | Sqh/Spdi<sup>c</sup> | *p*-value |
| --- | --- | --- | --- | --- | --- | --- | --- |
| WT/WT vs. WT/AA | 0.0029 | WT/EE vs. G915S/EE | 0.9973 | WT/WT vs. WT/AA | <0.0001 | WT/EE vs. G915S/EE | 0.7434 |
| WT/WT vs. WT/EE | <0.0001 | Q266P/AA vs. Q266P/EE | 0.8334 | WT/WT vs. WT/EE | <0.0001 | WT/EE vs. R931Q/AA | <0.0001 |
| WT/WT vs. Q266P/AA | <0.0001 | Q266P/AA vs. G915S/AA | <0.0001 | WT/WT vs. Q266P/AA | 0.9944 | WT/EE vs. R931Q/EE | <0.0001 |
| WT/WT vs. Q266P/EE | <0.0001 | Q266P/AA vs. G915S/EE | <0.0001 | WT/WT vs. Q266P/EE | 0.9808 | Q266P/AA vs. Q266P/EE | >0.9999 |
| WT/WT vs. G915S/AA | >0.9999 | Q266P/AA vs. R931Q/AA | 0.4937 | WT/WT vs. G915S/AA | 0.8118 | Q266P/AA vs. G915S/AA | 0.174 |
| WT/WT vs. G915S/EE | 0.0005 | Q266P/AA vs. R931Q/EE | <0.0001 | WT/WT vs. G915S/EE | 0.4961 | Q266P/AA vs. G915S/EE | 0.1137 |
| WT/WT vs. R931Q/AA | <0.0001 | Q266P/EE vs. G915S/AA | <0.0001 | WT/WT vs. R931Q/AA | 0.0043 | Q266P/AA vs. R931Q/AA | 0.0116 |
| WT/WT vs. R931Q/EE | <0.0001 | Q266P/EE vs. G915S/EE | <0.0001 | WT/WT vs. R931Q/EE | 0.9718 | Q266P/AA vs. R931Q/EE | >0.9999 |
| WT/AA vs. WT/EE | 0.0658 | Q266P/EE vs. R931Q/AA | 0.9976 | WT/AA vs. WT/EE | >0.9999 | Q266P/EE vs. G915S/AA | 0.1259 |
| WT/AA vs. Q266P/AA | <0.0001 | Q266P/EE vs. R931Q/EE | 0.0032 | WT/AA vs. Q266P/AA | <0.0001 | Q266P/EE vs. G915S/EE | 0.0861 |
| WT/AA vs. Q266P/EE | <0.0001 | G915S/AA vs. G915S/EE | 0.0007 | WT/AA vs. Q266P/EE | <0.0001 | Q266P/EE vs. R931Q/AA | 0.0282 |
| WT/AA vs. G915S/AA | 0.0062 | G915S/AA vs. R931Q/AA | <0.0001 | WT/AA vs. G915S/AA | 0.0052 | Q266P/EE vs. R931Q/EE | >0.9999 |
| WT/AA vs. G915S/EE | 0.411 | G915S/AA vs. R931Q/EE | <0.0001 | WT/AA vs. G915S/EE | 0.7439 | G915S/AA vs. G915S/EE | 0.9954 |
| WT/AA vs. R931Q/AA | <0.0001 | G915S/EE vs. R931Q/AA | <0.0001 | WT/AA vs. R931Q/AA | <0.0001 | G915S/AA vs. R931Q/AA | <0.0001 |
| WT/AA vs. R931Q/EE | <0.0001 | G915S/EE vs. R931Q/EE | <0.0001 | WT/AA vs. R931Q/EE | <0.0001 | G915S/AA vs. R931Q/EE | 0.1571 |
| WT/EE vs. Q266P/AA | <0.0001 | R931Q/AA vs. R931Q/EE | 0.1371 | WT/EE vs. Q266P/AA | <0.0001 | G915S/EE vs. R931Q/AA | <0.0001 |
| WT/EE vs.<br>Q266P/EE | <0.0001 | Q266P/AA vs.<br>Q266P/EE | 0.8334 | WT/EE vs.<br>Q266P/EE | <0.0001 | G915S/EE vs.<br>R931Q/EE | 0.0923 |
| WT/EE vs.<br>G915S/AA | <0.0001 | Q266P/AA vs.<br>G915S/AA | <0.0001 | WT/EE vs.<br>G915S/AA | 0.0048 | R931Q/AA vs.<br>R931Q/EE | 0.1161 |

<sup>a</sup>One-way ANOVA with Tukey's post-hoc analysis, N = 3, n = 11-93 cells.
<sup>b</sup>The overlap between Spdi (WT, Q266P, G915S, R931Q) and Sqh (WT, AA, EE).
<sup>c</sup>The overlap between Sqh (WT, AA, EE) and Spdi (WT, Q266P, G915S, R931Q).

