## Supplementary figures and images for "The *Drosophila* SPECC1L homolog, Split Discs, co-localizes with non-muscle myosin II and regulates focal adhesion dynamics"

### Supplemental Figure 1

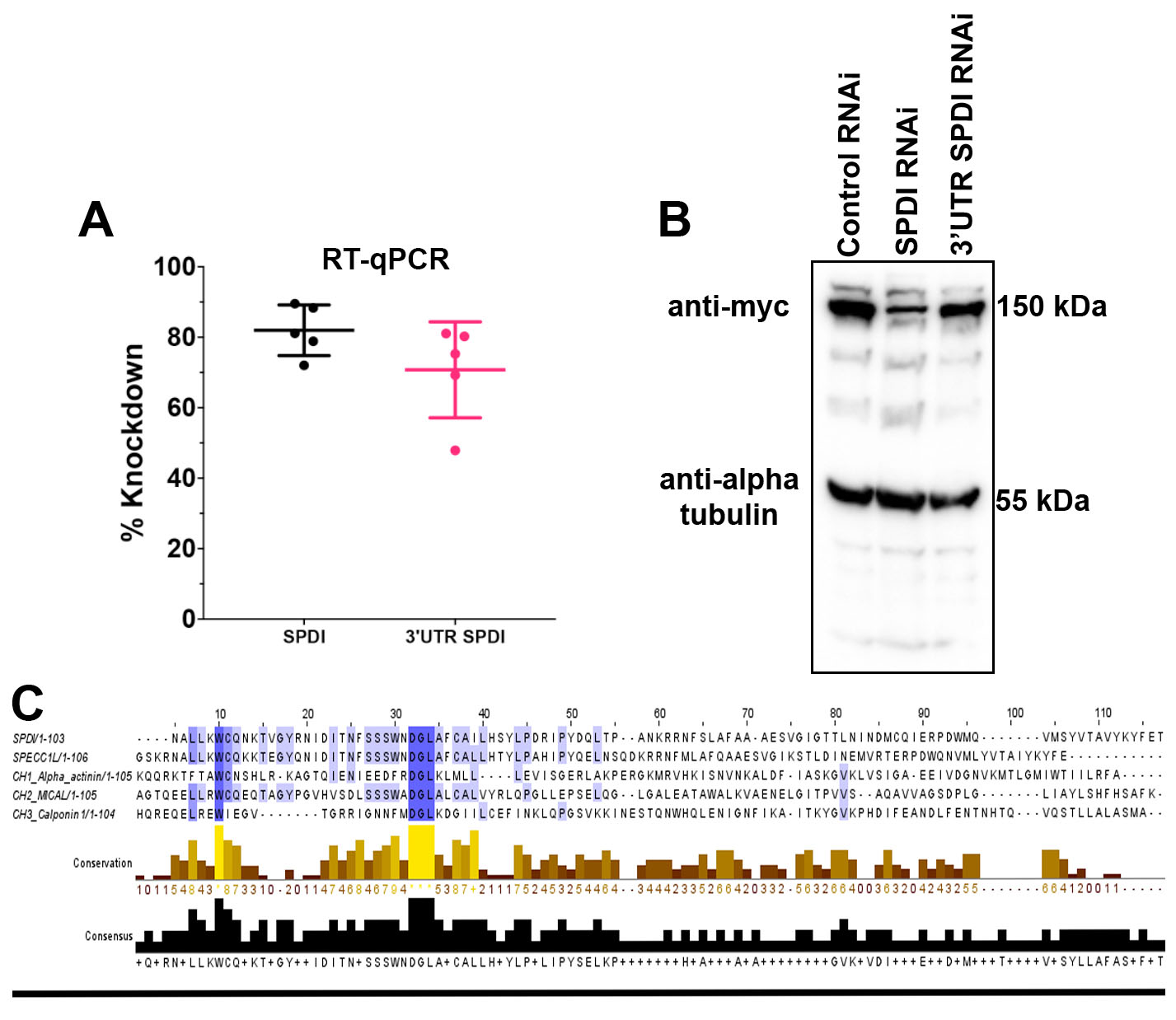

### Supplemental Figure 2

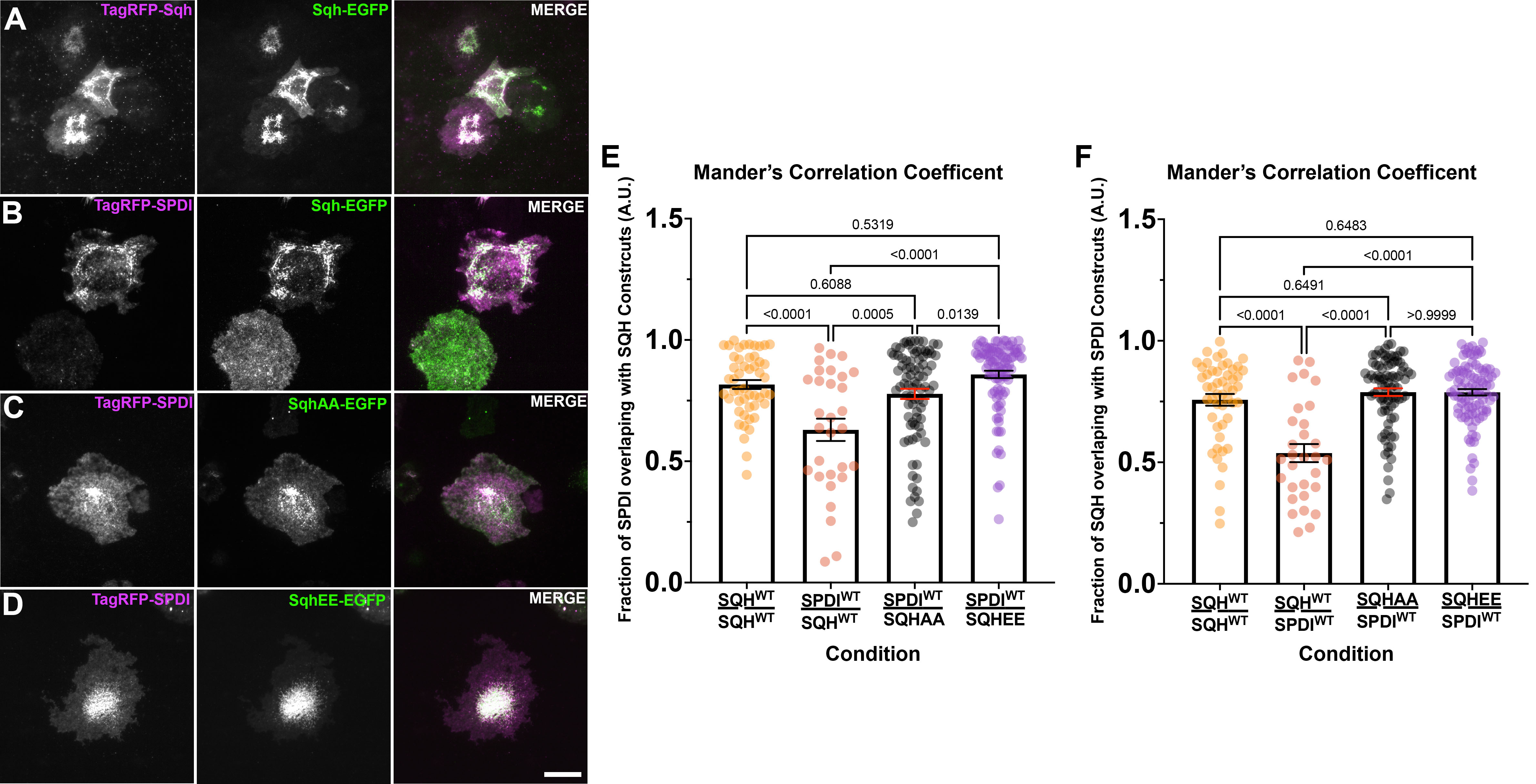

### Supplemental Figure 3

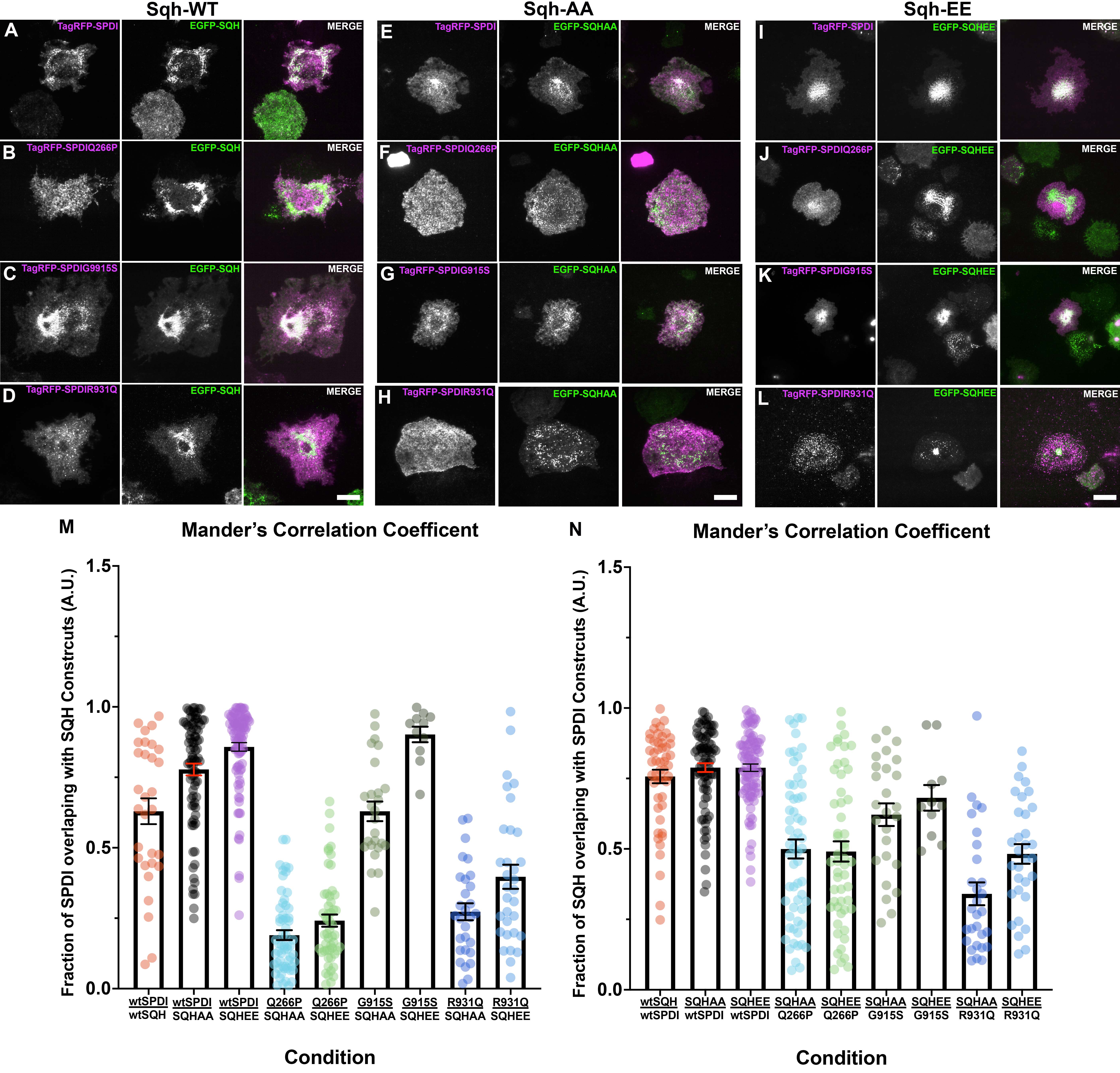

### Supplemental Figure 4

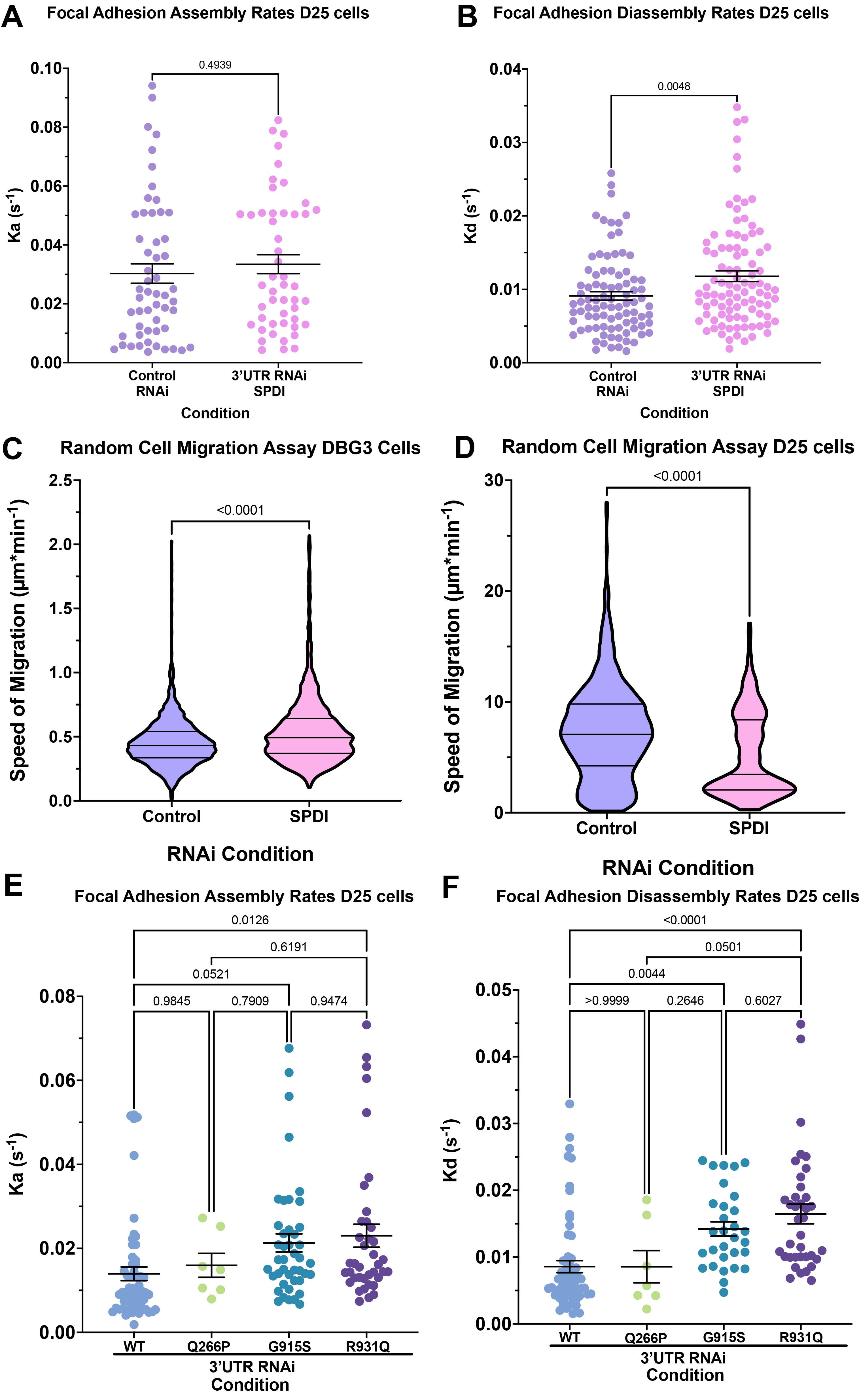
